# Spatial clustering of adhesion-deficient cells delays clearance and prolongs geometric unjamming signatures in epithelial tissues

**DOI:** 10.64898/2026.06.08.730803

**Authors:** Natalia Briñas-Pascual, Víctor Manso, Pilar Guerrero

**Affiliations:** Universidad Carlos III de Madrid, Departamento de Matemáticas, Grupo Interdisciplinar de Sistemas Complejos (GISC), Leganés, 28911, Madrid, Spain; ProCURE, Catalan Institute of Oncology, Oncobell, Bellvitge Institute for Biomedical Research (IDIBELL), 08908, L’Hospitalet del Llobregat, Barcelona, Spain; Universitat Autònoma de Barcelona, 08193, Bellaterra(Cerdanyola del Vallès), Barcelona, Spain

**Keywords:** Vertex Model, Unjamming Transition, E-cadherin, Cell Extrusion, Epithelial Topology, *T*_2_ removal event

## Abstract

Epithelial tissues maintain mechanical integrity through a balance between cell-cell adhesion and cortical contractility. Disruption of E-cadherin-mediated adhesion is a hallmark of epithelial-mesenchymal transition and cancer progression; yet how local adhesion defects propagate to tissue-scale mechanical changes remains poorly understood. Here, we use a two-dimensional vertex model, varying mutant cell fraction, spatial arrangement, and initial tissue disorder, to investigate how adhesion-deficient cells regulate epithelial mechanics.

We show that increasing mutant fraction drives the tissue towards geometric signatures associated with reduced mechanical rigidity, characterised by elevated cellular shape index and increased prevalence of non-hexagonal cells. Crucially, spatial organisation modulates tissue mechanics independently of mutant fraction. For identical mutant fractions, randomly distributed mutants undergo rapid, mutually independent T_2_-mediated removal, producing only transient shape-index perturbations. Clustered mutants, by contrast, undergo sequential boundary removal, delaying elimination and sustaining elevated shape index in the surrounding tissue; this persistent elevation induces topological disorder that outlasts mutant clearance itself. We show that these signatures arise from the reduced heterotypic interface exposure of interior cluster cells, unifying clearance kinetics, shape-index elevation, and topological disorder as projections of a single mechanism, with implications for understanding early-stage cancer progression.

## 1. Introduction

Epithelial tissues function as dynamic physical barriers that delineate internal biological compartments from their external environment. To sustain this barrier, epithelial monolayers must maintain robust mechanical integrity while preserving the plasticity required for morphogenesis, wound healing, and cellular turnover [1, 2, 3, 4, 5]. This balance is governed by the mechanical interplay between cell–cell adhesion, and actomyosin-generated cortical tension, which together determine effective interfacial tensions and regulate junctional remodelling and cell rearrangements [6, 7, 8]. In epithelial tissues, cell–cell adhesion is primarily mediated by cadherin complexes, including E-cadherin, and contractile forces are generated by the actomyosin cytoskeleton [1, 9, 8]. Under homeostatic conditions, this coupling helps tissues remain in a jammed, solid-like state, in which large-scale topological rearrangements are energetically suppressed [10, 11, 12].

Loss of E-cadherin is a defining feature of both physiological development and pathological progression, most notably during epithelial-to-mesenchymal transition (EMT) and cancer metastasis [13, 14, 15, 16, 17, 18]. When junctional adhesion is compromised, the force balance at cell-cell interfaces is disrupted, often triggering cell extrusion [19, 20, 21]. While homeostatic extrusion typically eliminates apoptotic or overcrowded cells [19], aberrant extrusion associated with adhesion defects may facilitate invasive behaviour and the dissemination of tumorigenic cells [22, 23, 24, 25, 26].

Mechanical rigidity and jamming–unjamming transitions provide a useful framework for understanding collective epithelial rearrangements [11, 12, 10]. The geometric packing and shape of epithelial cells are determined by the competition between cortical tension and adhesion [10, 7]. In vertex-based descriptions, this transition is commonly quantified by the cellular shape index, *p* = *P/ √A*, where *P* is the cell perimeter and *A* is the cell area. This dimensionless order parameter has a critical threshold, *p*^∗^ 3.81, that marks the transition from a jammed solid-like state to an unjammed, fluid-like regime [27, 28, 11, 29, 30]. Because this threshold was derived for homogeneous systems, in the heterogeneous tissues considered here we use it as a reference for unjamming-associated geometric changes rather than as a direct measurement of mechanical rigidity.

Although jamming–unjamming transitions have been extensively characterised in homogeneous tissues [31, 32, 33, 34, 35, 36], real pathological tissues are inherently heterogeneous [37, 38]. More recent work has shown that such heterogeneity can itself modify collective tissue mechanics. In vertex models, heterogeneity in preferred cell shape can alter the tissue shear modulus and rigidity transition [39], higher-order vertex structures and multicellular rosettes can regulate fluid–solid transitions [30], and spatial correlations in preferred cell shape or target area can shift the rigidity transition [40]. Thus, both cell-level mechanical heterogeneity and its spatial organisation are established determinants of collective tissue behaviour. Experimental studies have likewise shown that tumours exhibit spatial heterogeneity in their mechanical properties, which can influence cancer invasion and progression [41, 42, 43, 44, 39, 45].

The spatial context of mechanically distinct cells, such as defective cells in early tumours, is particularly relevant in epithelial tissues. These defective cells rarely appear as isolated defects. Instead, they often emerge through clonal expansion, forming spatially correlated clusters within an otherwise wild–type epithelium [46, 47, 17, 22, 23]. Previous studies have shown that the fate of defective epithelial cells depends on tissue architecture and properties of the surrounding population [1, 12, 48, 49]. In particular, Ras-transformed epithelial cells can undergo apical extrusion when surrounded by normal epithelial cells, whereas this behaviour is strongly altered when transformed cells are surrounded by other transformed cells [50]. These observations motivate considering not only the abundance of mechanically defective cells, but also their exposure to heterotypic contacts with the surrounding wild-type tissue.

A substantial body of work on differential adhesion and interfacial tension further demonstrates that interfaces between mechanically distinct cell populations can play an active role in tissue organisation. Differential interfacial tension can drive cell sorting, segregation, and tissue rearrangement [6, 7, 8], while elevated tension specifically at heterotypic interfaces can maintain boundaries between distinct cell populations [51]. In confluent epithelia, vertex-model and experimental studies have also demonstrated small-scale demixing in mixtures of wild-type and E-cadherin-deficient keratinocytes [52]. More recently, heterotypic interfacial tension between wild-type and oncogenic populations has been identified as an important determinant of oncogenic clone behaviour [53].

Here, we address a more specific question: for a fixed fraction of adhesion-deficient cells, differing from wild-type cells only in their interfacial line tension, how does their spatial arrangement alter their heterotypic exposure to the surrounding wild-type tissue, and how does this in turn influence T_2_-mediated clearance and the subsequent geometric and topological response of the epithelium? Here, adhesion-deficient cells, referred to throughout as mutant cells, are introduced either as randomly scattered defects or as spatially contiguous clusters, while the initial degree of tissue disorder is independently controlled.

We hypothesise that the spatial organisation of adhesion-deficient cells regulates their heterotypic exposure to the surrounding wild-type tissue and thereby modulates T_2_-mediated clearance dynamics and the duration of the resulting geometric and topological perturbations. Specifically, we investigate how mutant spatial arrangement and initial structural disorder affect mutant-cell clearance kinetics, local and global shape-index dynamics, and the lifetime of extrusion-induced topological disorder.

To address these questions, we use a two-dimensional vertex model [1, 2], in which adhesion-deficient cells are introduced either as randomly scattered defects or as compact clonal clusters. We report four results. First, spatial clustering delays T_2_-mediated clearance across the entire parameter space explored, establishing spatial arrangement as a control parameter alongside mutant density. Second, this delay follows from the heterotypic exposure per surviving mutant cell, which in clustered configurations begins low and rises as the clone is eroded, following the scaling expected for a compact two-dimensional domain; notably, the cells that survive longest are those carrying the highest absolute effective edge tension in the tissue, so it is the contrast in effective tension across the heterotypic interface, and not its magnitude, that drives removal. Third, the resulting prolonged mechanical loading of the surrounding wild-type tissue sustains an elevated shape index in the neighbouring cells long after random configurations have relaxed, and leaves a topological imprint, a broad shift of the local polygon distribution towards classes away from six, that persists beyond clearance itself. Fourth, this imprint has a direct mechanical correlate: the local effective edge tension of the wild-type cells that surrounded the clone remains elevated once the clone is gone. Together, these findings identify the spatial organisation of adhesion-deficient cells as a structural control variable of the geometric signatures of epithelial unjamming, acting through heterotypic interface exposure, with implications for understanding early-stage cancer progression.

## 2. Mathematical Framework

We model the mechanics of epithelial tissues using a two-dimensional vertex model [1, 54, 2]. In this framework, the apical surface of the epithelium is represented as a polygonal tessellation whose vertices correspond to tricel-lular junctions. Cells share edges (junctions) and meet at vertices, forming a dynamic network that evolves in response to mechanical forces. The mechanical state of the tissue is determined by minimising a potential energy functional that captures the competition between cell elasticity, cortical contractility, and cell-cell adhesion.

### 2.1. Energy functional

Following [1], the total mechanical energy of a tissue composed of *N* cells is

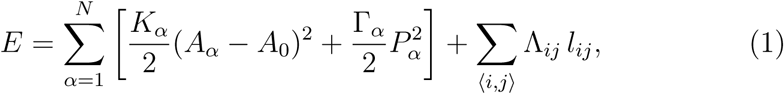

where *A_α_* and *P_α_* denote the area and perimeter of cell *α*, respectively. The first term represents area elasticity, penalising deviations from a preferred area *A*_0_ and reflecting the effective incompressibility of epithelial cells. The second term describes cortical contractility arising from the actomyosin cortex, with Γ*_α_* controlling its strength. The third term accounts for interfacial tension along cell-cell junctions, where *l_ij_* is the length of the edge connecting vertices *i* and *j*, and Λ*_ij_* is the effective line tension combining cortical tension and cadherin-mediated adhesion [34].

### 2.2. Adhesion loss and mechanical heterogeneity

In homeostatic tissues, mechanical parameters are typically assumed uniform across the lattice. To model the loss of E-cadherin-mediated adhesion, we introduce mechanical heterogeneity by assigning interface-dependent values of Λ*_ij_* [28, 33]:

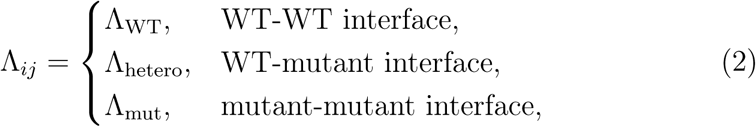

where, following [23], the heterotypic value is given by the arithmetic mixing rule

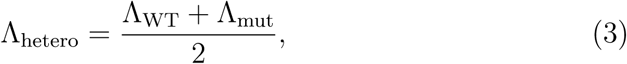

with Λ_mut_ *>* Λ_WT_ [1, 23]. Here, Λ*_ij_* represents an effective junctional line tension incorporating the balance between cortical tension and cell–cell adhesion, and we model adhesion deficiency by increasing this effective parameter at interfaces involving mutant cells. For the parameter values used here, Λ_WT_ = 0.075 and Λ_mut_ = 0.68, giving Λ_hetero_ = 0.3775.

The line tension of an edge is therefore determined locally, from the identities of the two cells sharing it, and is re-evaluated whenever the local adjacency changes through a topological rearrangement, so that newly created interfaces receive the value of Λ*_ij_* corresponding to their new neighbours. Mutant identity itself is assigned before the mechanical simulation begins and remains fixed until the corresponding cell is removed through a T_2_ event; it does not spread dynamically. The spatial distribution of line tensions thus evolves only through changes in tissue topology and through mutant removal, rather than through conversion of WT cells into mutants.

All other mechanical parameters, including area elasticity *K*, cortical contractility Γ, and preferred area *A*_0_, are identical for WT and mutant cells. Mechanical heterogeneity is therefore introduced exclusively through the interface-dependent line-tension coefficient Λ*_ij_*, and any difference in the mechanical state of wild-type and mutant cells is a consequence of the line-tension contrast alone rather than of an additional.

To investigate the role of spatial organisation, mutant cells are introduced either as randomly distributed defects across the lattice or as contiguous clusters occupying a compact region of the tissue, mimicking clonal expansion observed in E-cadherin-related pathologies [23]. The fraction of mutant cells in the tissue is defined as *f*_mut_ = *N*_mut_*/N*_cells_, where *N*_mut_ is the number of adhesion-deficient cells and *N*_cells_ is the total number of cells, and is varied between 0 and 0.30 in increments of 0.025.

### 2.3. Initial tissue disorder

Biological epithelial tissues rarely display perfectly regular cellular packing. Even under homeostatic conditions, monolayers exhibit geometrical disorder due to growth, mechanical fluctuations, and cell turnover; in pathological conditions such as cancer, this disorder is often amplified [14, 12, 34, 1].

To reproduce this variability, simulations are initialised from a regular hexagonal lattice whose node positions are perturbed as

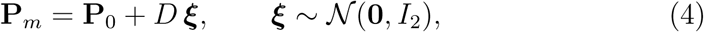

where **P**_0_ are the unperturbed node coordinates, *D* is a distortion parameter controlling the perturbation amplitude, and ***ξ*** is a random vector drawn from a bivariate normal distribution with zero mean and identity covariance. The perturbation amplitude *D* is expressed in units of the hexagonal lattice spacing (i.e. the distance between nearest-neighbour vertices), so that *D* = 0 corresponds to a perfectly regular hexagonal lattice, while larger values introduce increasing structural disorder. In this study, *D* is varied between 0 and 0.7 in increments of 0.05. Simulations use periodic boundary conditions throughout.

### 2.4. Shape index and geometric signatures of unjamming

Geometric signatures of unjamming in epithelial vertex models are commonly characterised through the *cellular shape index*:

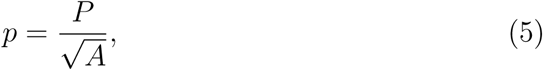

a dimensionless quantity that acts as an order parameter for jamming transitions in epithelial tissues [11, 12, 10]. The minimum possible value for the shape index in a planar tessellation is *p* ≈ 3.72, attained for the maximally ordered configuration of regular hexagons [55]; thus, 3.72 represents the strict lower bound of *p*. When *p* exceeds a critical threshold (*p*^∗^ ≈ 3.81), energy barriers for neighbour exchanges vanish and the tissue displays geometric signatures associated with reduced mechanical rigidity, consistent with proximity to an unjamming-like threshold [28, 11, 12]. This threshold was derived for homogeneous tissues; because it was derived for homogeneous systems, we use it here as a reference for unjamming-associated geometric changes rather than as a direct measurement of mechanical rigidity, and its application to heterogeneous systems such as ours is discussed further in Section 3.4. In our simulations, ❬*p*❭ is monitored at two spatial scales. The global mean shape index, ❬*p*❭ _global_, is averaged over all cells at each time point. The local mean shape index, ❬*p*❭ _local_, is averaged over the wild-type cells in direct contact with a mutant cell (or, in the case of clustered arrangements, with the boundary of the cluster). This neighbourhood set, referred to throughout as the mutant-neighbourhood wild-type cells, is dynamically updated throughout the lifespan of each mutant cell, and is fixed once that mutant cell is cleared. The local signal therefore continues to report on the same tissue region after mutant clearance, capturing the persistent mechanical and topological imprint left where the adhesion-deficient cells were located. The same local/global distinction is applied to topological measurements of polygon class frequencies and hexagonal packing deviation.

### 2.5. Heterotypic interface exposure and effective edge tension

To characterise the mechanical consequences of heterotypic WT-mutant contacts, we introduce two complementary quantities describing the exposure of mutant cells to the surrounding wild-type tissue and the mechanical loading this exposure generates.

The first is the heterotypic interface length, *L*_het_(*t*), defined as the summed length of all edges connecting a wild-type cell to a mutant cell at time *t*. Normalising by the number of surviving mutant cells, *N*_mut_(*t*), defines the heterotypic interface exposure per surviving mutant cell,

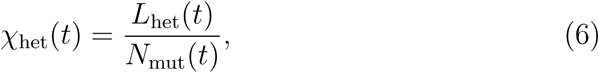

a purely geometric measure of the exposure of the surviving mutant population to wild-type neighbours. Separating *χ*_het_ from *L*_het_ distinguishes a reduction in interface caused by the loss of mutant cells from a reduction in interface per surviving mutant cell.

The second quantity is mechanical. While the line tension Λ*_ij_* in equation (1) sets the explicit interfacial contribution, the force acting on an edge also receives a contribution from the cortical contractility of each of the two cells sharing it, since *∂P_α_/∂l_ij_* = 1 for any edge belonging to cell *α*. Differentiating the total energy with respect to the length of edge *ij*, shared by cells *α* and *α*^r^, at fixed geometry, gives the effective edge tension

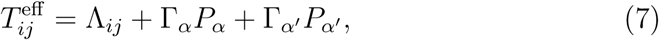

the standard quantity used in vertex-model stability analyses [1, 28]. An edge with *T_ij_^eff^ >* 0 favours contraction (and thus *T*_1_/*T*_2_ transitions), while *T_ij_^eff^ <* 0 favours expansion; the condition *T_ij_^eff^* = 0 corresponds to the marginal edge stability criterion of [28]. For each stored snapshot, *T* ^eff^ is computed for every edge using the corresponding line tension Λ*_ij_* ∈ {Λ_WT_, Λ_hetero_, Λ_mut_} (equation (2)) and the instantaneous perimeters *P_α_*, *P_α_′* of the two cells sharing the edge. Averaging *T_ij_^eff^* over the edges of a given cell defines a per-cell effective tension,

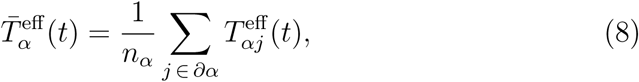

where *∂α* denotes the set of edges bounding cell *α* and *n_α_* its number of neighbours.

Finally, to assess whether mutant clearance leaves a lasting mechanical signature in the surrounding tissue, we apply the same local/global decomposition used for the shape index (Section 2.4) to the effective edge tension of wild-type edges, defining ⟨*T* ^eff^⟩_local_ over the edges of the mutant-neighbourhood wild-type cells, tracked throughout the simulation including after mutant clearance, and ⟨*T* ^eff^ ⟩_global_ over all edges in the tissue.

### 2.6. Vertex dynamics and topological transitions

Tissue dynamics are simulated assuming an over-damped motion appropriate for epithelial monolayers interacting with a substrate [2]. The position of each vertex *i* evolves as

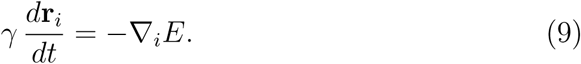

Here, *γ* denotes the homogeneous vertex friction coefficient and sets the mechanical relaxation timescale of the tissue. For a constant friction coefficient, the time variable can be rescaled as *τ* = *t/γ*, such that changes in *γ* primarily modify the rate at which the tissue relaxes rather than the underlying energy landscape [56].

We set *γ* = 0.02 to obtain a convenient temporal parametrisation in which vertex relaxation and topological rearrangements can be readily resolved within a computationally manageable simulation window. This value is therefore used to set the simulation timescale and should not be interpreted as an experimentally calibrated estimate of epithelial friction. This rescaling is appropriate here because the model does not include an additional independently prescribed biological timescale, such as cell-cycle progression or proliferation.

Two types of discrete topological rearrangements are permitted:

#### T_1_ transitions (cell intercalation)

When the length of a junction *l_ij_* falls below a threshold *ɛ_T_*_1_ = 0.04 (in units of *A*_0_), the edge is removed and replaced by a new edge connecting the previously opposite vertices, changing the neighbour relationships between cells [28].

#### T_2_ transitions (cell extrusion)

Cell extrusion is not imposed through an independent area criterion but arises as a topological consequence of successive *T*_1_ neighbour exchanges. As a cell repeatedly loses edges through *T*_1_ events its polygon class decreases, and once a *T*_1_ event would leave it bounded by only two edges the cell is removed from the tissue and the surrounding connectivity is updated to maintain a consistent mesh. Cell elimination is therefore governed entirely by *ɛ_T_*_1_, which is the only topological parameter of the model; no separate extrusion threshold is prescribed. For reference, the smallest cell that can persist is a triangle whose edges are all at the *T*_1_ threshold length, so that cells disappear at an area scale of order 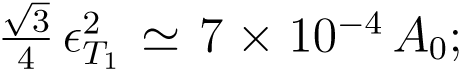 this is a consequence of the dynamics rather than a criterion applied to them. This provides a geometric representation of extrusion-like events in epithelial monolayers [19, 20]. We note that this geometric description represents a simplified account of biological extrusion, which involves active actomyosin contraction and apoptotic signalling not explicitly modelled here [21, 24].

Equations of motion are integrated using a forward Euler scheme with time step *dt* = 0.001. Simulations were performed using the parameter values listed in table 1.

**Table 1:**
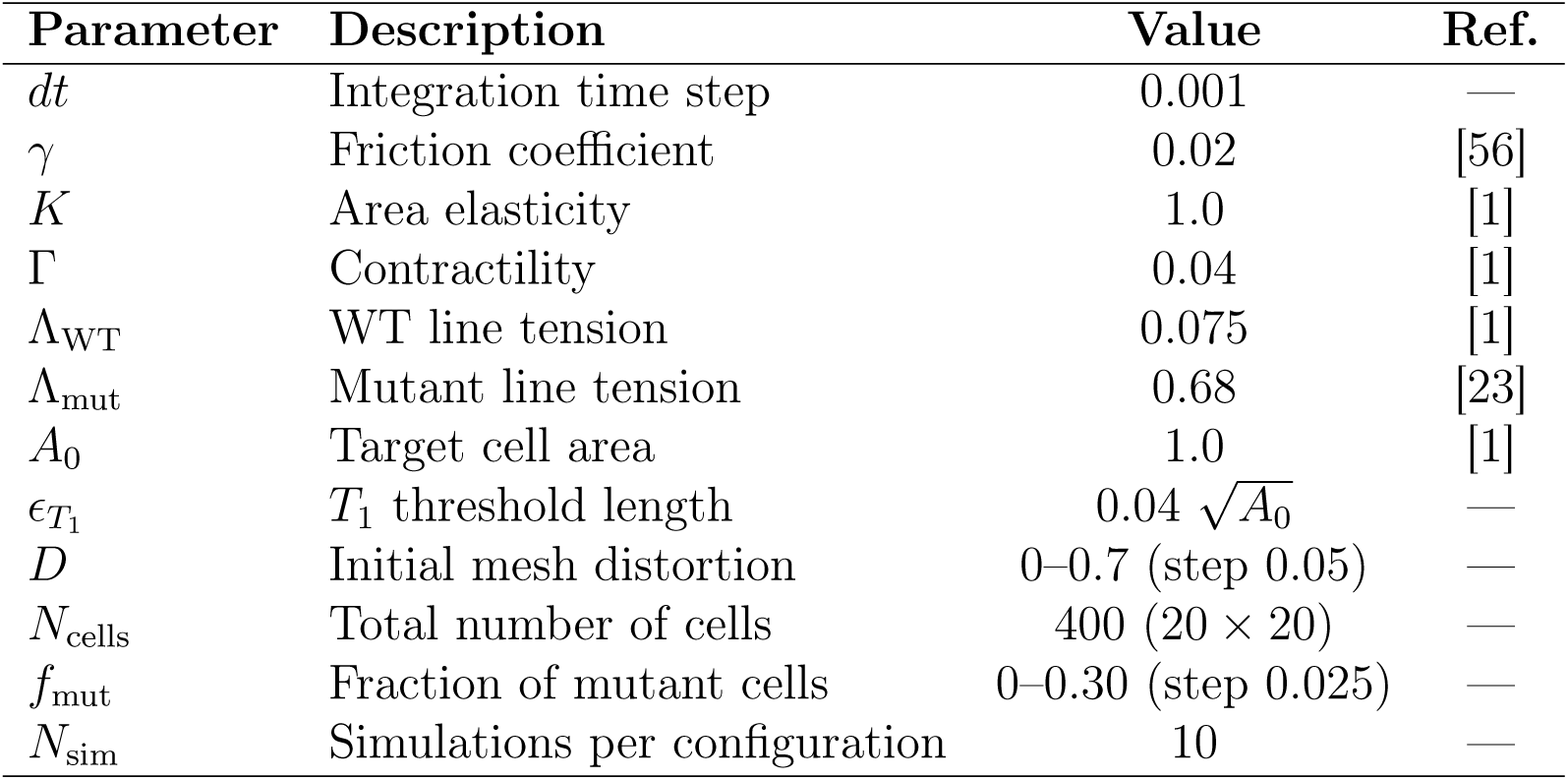
Parameters used in vertex model simulations.

### 2.7. Spatial arrangement of adhesion-deficient mutant cells

The clustered configuration is generated entirely at *t* = 0; mutant identity does not spread dynamically during the mechanical simulation. The cluster of mutant cells is initialised by selecting a single seed cell uniformly at random from the tissue. The cluster is then grown iteratively based on topological adjacency. Specifically, the algorithm first identifies the set of first-order neighbours of the seed cell and randomly selects from this set to assign mutant identities. Once this initial neighbour set is exhausted, the neighbourhood is recomputed as the set of cells adjacent to any cell currently in the mutant cluster, effectively expanding the cluster shell by shell in terms of graph distance. This process is repeated until the desired number of mutant cells, [*f*_mut_ *N*_cells_], is reached or no further expansion is possible. The procedure described above is solely an initialization algorithm for constructing a connected mutant population of prescribed size. Once the desired mutant fraction has been assigned, mutant identities remain fixed, and subsequent changes in mutant number arise only through T_2_-mediated removal.

This construction mimics clonal expansion in tumour growth, where a single progenitor gives rise to a spatially contiguous population. It ensures that the mutant cluster remains connected and localised. Because growth proceeds through successive neighbour shells, the resulting shape tends towards radial symmetry in highly ordered tissues, while remaining more irregular in disordered environments. Consequently, cluster compactness emerges from the interplay between lattice structure and random growth, rather than being uniquely determined by *f*_mut_. For comparison, in random configurations, mutant cells are selected uniformly at random without replacement from the full lattice.

## 3. Results

Before comparing the late-time mechanical and topological properties of the tissues, we first examined the evolution of the total mechanical energy. In both random and clustered configurations, the energy decreases markedly during the initial transient and subsequently approaches a stable plateau across the range of mutant fractions and initial mesh distortions considered (Supplementary Figure S1). This behaviour indicates that the tissues progressively relax towards mechanically stationary configurations. Although the relaxation timescale differs between random and clustered arrangements, the energy has plateaued in both cases well before *t* = 15000, and we therefore evaluate all post-extrusion observables at that single, common time. The geometric, mechanical and topological observables discussed below were thus evaluated once the initial relaxation dynamics had effectively ceased and at the same time point for both configurations, allowing differences between random and clustered tissues to be interpreted as differences between their relaxed states rather than as differences in relaxation rate or in observation time.

### 3.1. Adhesion-deficient mutant cells perturb epithelial packing

We first investigate how the spatial distribution of adhesion-deficient mutant cells alters the geometric organisation and mechanical stability of epithelial tissues. Simulations were initialised from originally hexagonal lattices, distorted through controlled levels of structural noise *D* and varying fractions of mutant cells. Adhesion loss was implemented by assigning increased line tensions to interfaces involving mutant cells, following equation (2), reflecting the mechanical consequence of reduced E-cadherin-mediated adhesion [1, 14]. The parameters used are described in table 1.

Figure 1 illustrates the morphological evolution, at three different times, of tissues containing randomly distributed (panel A) and clustered (panel B) populations of adhesion-deficient cells (*D* = 0.25, *f*_mut_= 0.25) and the dynamics are illustrated in electronic supplementary material, videos S1 and S2, table 2. In both cases, the line tension imbalance at mutant interfaces drives a sequence of *T*_2_-mediated cell removal events that progressively eliminate mutant cells in the vertex model [19, 20]. However, the spatial arrangement of defects markedly affects how this process unfolds: randomly distributed mutants undergo T_2_-mediated removal rapidly and independently, whereas clustered mutants generate coordinated shape-index elevations at the cluster periphery (quantified in figure 5) that persist in the surrounding wild-type matrix for longer periods.

**Figure 1:**
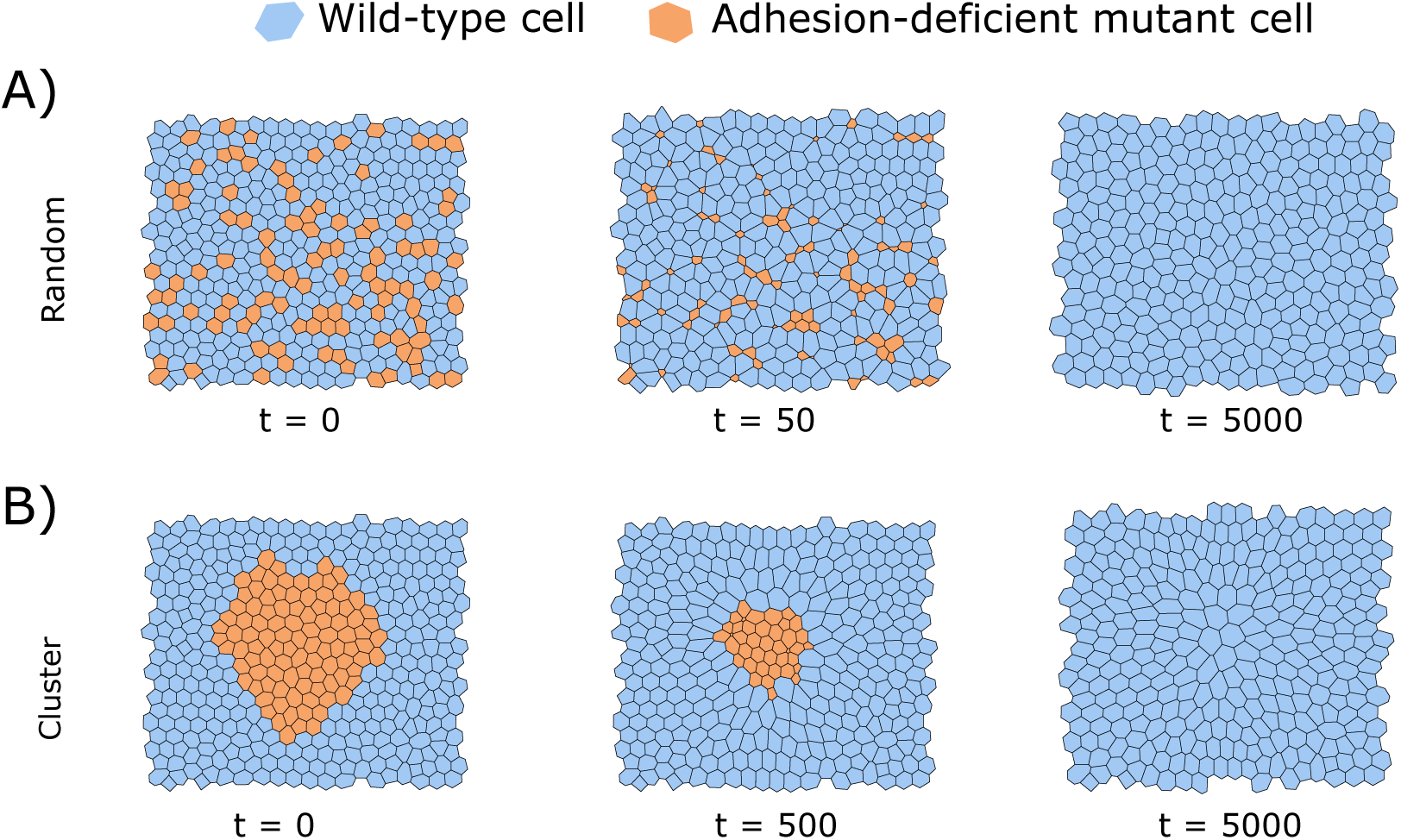
Time evolution of epithelial tissues containing adhesion-deficient cells (*D* = 0.25, *f*_mut_ = 0.25) distributed randomly (A, at *t* = 0, 50, and 5000) or as a contiguous cluster (B, at *t* = 0, 500, and 5000); the different time points reflect the substantially slower extrusion dynamics of clustered configurations. Mutant cells are shown in orange and wild-type cells in blue. The rest of the simulation parameters are described in table 1.

### 3.2. Clustering slows T_2_-mediated adhesion-deficient mutant removal

Before analysing how shape-index elevations spread through the tissue, we first quantify a fundamental difference between the two spatial configurations: the kinetics of mutant cell removal.

**Table 2:**
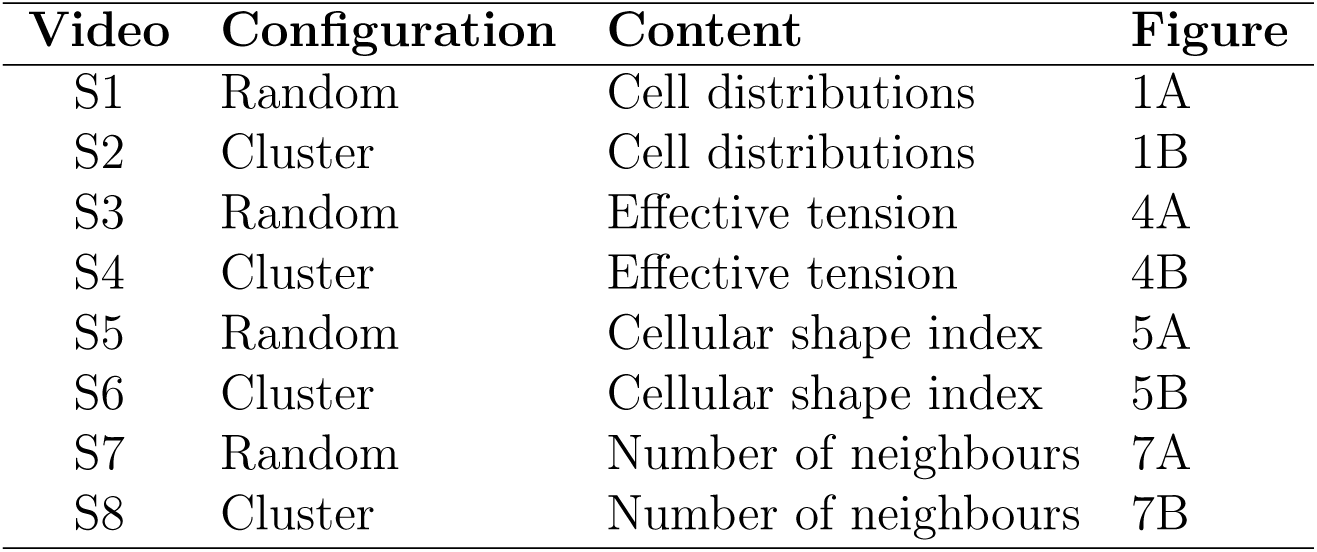
Electronic supplementary material videos (*D* = 0.25, *f*_mut_ = 0.25)

| Video | Configuration | Content | Figure |
| --- | --- | --- | --- |
| S1 | Random | Cell distributions | 1A |
| S2 | Cluster | Cell distributions | 1B |
| S3 | Random | Effective tension | 4A |
| S4 | Cluster | Effective tension | 4B |
| S5 | Random | Cellular shape index | 5A |
| S6 | Cluster | Cellular shape index | 5B |
| S7 | Random | Number of neighbours | 7A |
| S8 | Cluster | Number of neighbours | 7B |

To characterise this process, we define the extrusion time *t_x_*_%_ as the earliest time at which the fraction of remaining adhesion-deficient cells falls below a given threshold *x*,

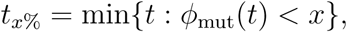

where *φ*_mut_(*t*) = *N*_mut_(*t*)*/N*_mut_(0). This measure allows us to compare the timescale of mutant-cell clearance across spatial configurations and across different stages of the extrusion process. Figure 2 illustrates the temporal evolution of *φ*_mut_ for random and clustered initial distributions, considering two levels of initial disorder *D* (0.1 and 0.5) and the full range of mutant fractions *f*_mut_ (from 0 to 0.30 in steps of 0.025, encoded by colour intensity).

**Figure 2:**
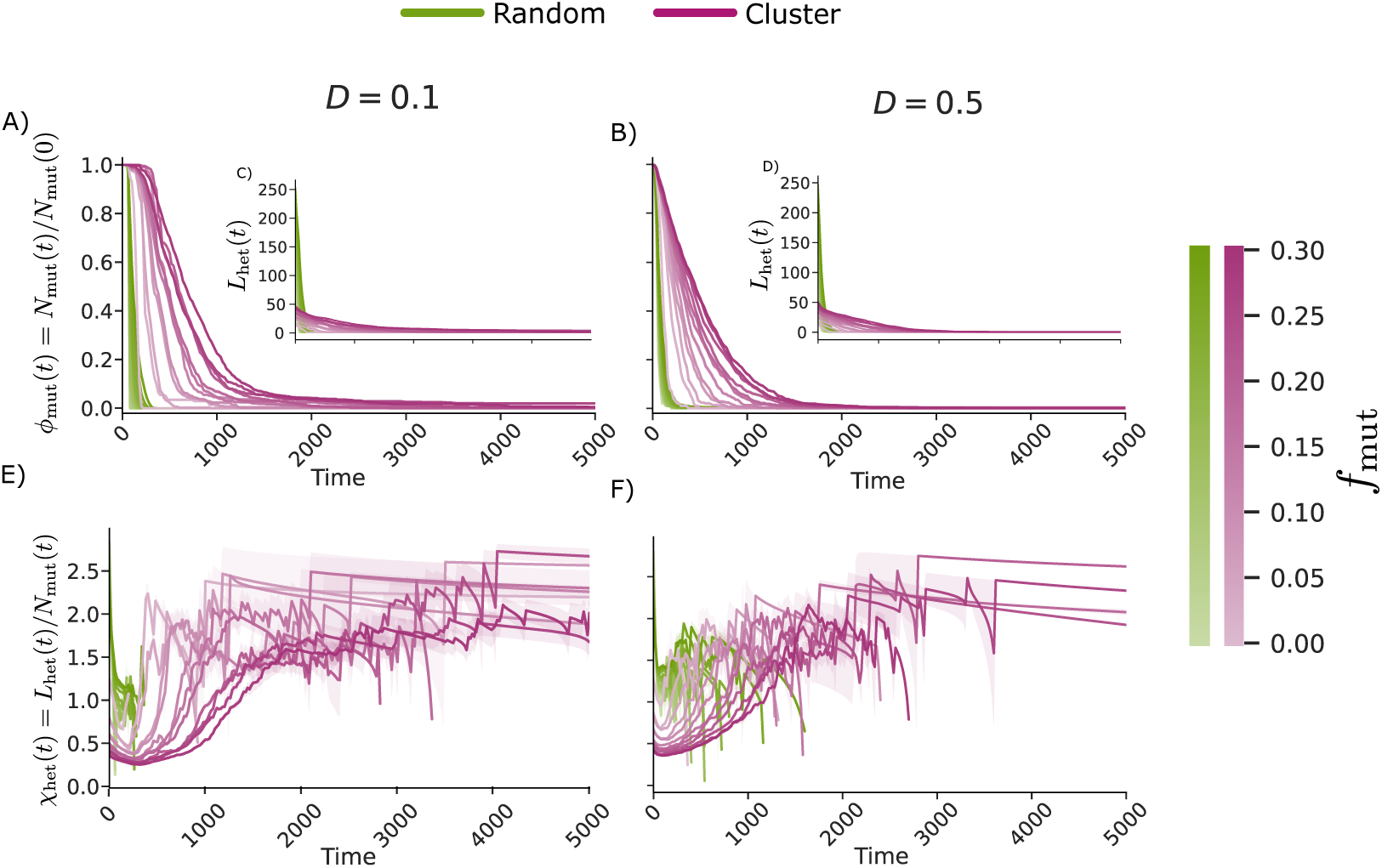
Mutant clearance kinetics and heterotypic interface exposure for randomly distributed (green) and clustered (magenta) adhesion-deficient cells, at low (*D* = 0.1, left column) and high (*D* = 0.5, right column) initial mesh distortion. (A, B) Temporal evolution of the fraction of remaining mutant cells, *φ*_mut_(*t*) = *N*_mut_(*t*)*/N*_mut_(0). (C, D) Insets: total heterotypic interface length *L*_het_(*t*), the summed length of all wild-type–mutant edges. (E, F) Heterotypic interface exposure per surviving mutant cell, *χ*_het_(*t*) = *L*_het_(*t*)*/N*_mut_(*t*). Colour intensity encodes the initial mutant fraction, from *f*_mut_ = 0.025 (light) to *f*_mut_ = 0.30 (dark). Each curve represents the mean over *N*_sim_ = 10 independent simulations; shaded bands indicate the standard error of the mean (SEM). Simulations were run to *t* = 15000; the time axis is truncated at *t* = 5000 to resolve the extrusion dynamics. At late times *χ*_het_ is evaluated over the few mutant cells that remain, which accounts for the increasingly stepped appearance of the clustered curves in (E, F). The rest of the simulation parameters are described in table 1.

The shaded bands around each mean curve represent the standard error of the mean (SEM) across *N*_sim_ = 10 independent realisations. The consistently narrow SEM bands suggest that extrusion dynamics are highly reproducible for fixed parameters, and that the observed differences between spatial configurations substantially exceed the inter-simulation variability throughout the explored parameter space.

In all simulations, the fraction of adhesion-deficient cells decreases monotonically. However, the rate of this process depends strongly on the spatial organisation of the mutants. In random configurations (figure 2, green lines), mutant cells are dispersed throughout the tissue and are typically surrounded by wild-type (WT) neighbours. This arrangement maximises the number of heterotypic interfaces, where the contrast in line tension (Λ_hetero_ Λ_WT_) is largest. As a result, isolated mutant cells experience strong local mechanical imbalance and undergo T_2_-mediated removal rapidly and largely independently of one another [23]. Extrusion events therefore occur in parallel across the tissue, producing a relatively fast global elimination of defects.

In contrast, clustered configurations exhibit significantly slower extrusion dynamics than random ones. Mutant cells located in the interior of the cluster are surrounded primarily by other mutant cells and therefore share homotypic Λ_mut_ interfaces. This configuration reduces the number of heterotypic WT-mutant interfaces experienced by interior cells, limiting the mechanical contrast that drives T_2_-mediated removal. The delayed elimination of clustered mutants should, therefore, be interpreted primarily as a consequence of reduced heterotypic exposure of interior cells, rather than as a reduction of their absolute interfacial tension. Consequently, T_2_-mediated removal proceeds sequentially from the cluster boundary towards its interior, imposing an outside-in elimination mechanism. This geometric constraint serialises the extrusion process and substantially delays the removal of the mutant population.

The ratio *t*_50%_*^cluster^*/*t*_50%_*^random^*, shown in Supplementary Figure S2 (right column, top row), quantifies this delay across the full parameter space. Clustered configurations consistently satisfy *t*_50%_*^cluster^* > *t*_50%_*^random^* with the ratio exceeding 2 throughout the entire (*D, f*_mut_) parameter space, confirming that spatial aggregation robustly hinders mutant clearance under all tested conditions. The ratio grows monotonically with *f*_mut_, and does so at both low and high initial disorder: it is smallest for the smallest clones, which comprise only a handful of cells and are therefore barely distinguishable from a dispersed configuration, and largest at the highest mutant fractions, where the shielded interior of the clone is largest relative to its boundary. The delay conferred by clustering therefore scales with clone size, as the exposure argument of Section 3.3 predicts.

Importantly, this kinetic difference persists across the explored parameter space. In both spatial configurations, increasing *f*_mut_ prolongs mutant persistence, as reflected by larger values of *t*_50%_ and *t*_5%_ (Supplementary Figure S2). Higher initial tissue distortion *D* tends to accelerate the early stages of extrusion in both configurations, consistent with the lower energy barriers to topological rearrangement in tissues that start further from regular hexagonal packing [28, 34].

However, the random and clustered configurations diverge markedly at late stages of extrusion, as captured by *t*_5%_. In random configurations, a striking inversion emerges: while higher distortion accelerates early extrusion, complete clearance of the last remaining mutants is achieved faster in tissues with lower initial disorder. This behaviour is consistent with the progressive isolation of randomly distributed mutants, whose final elimination becomes more efficient in ordered tissues once the majority of the mutant population has already been removed.

By contrast, clustered mutants do not display this inversion in behaviour. Instead, lower initial disorder consistently prolongs mutant persistence throughout the entire extrusion process, and the delay associated with clustering remains robust under all tested conditions. These results demonstrate that spatial organisation acts as an independent control parameter governing the timescale of tissue remodelling, allowing adhesion-deficient cells to persist within the epithelium for substantially longer periods when arranged in clusters. The narrow SEM bands in figure 2 suggest that these differences are unlikely to be artefacts of stochastic variability: the systematic separation between random and clustered curves exceeds the inter-simulation spread across the entire parameter space explored, supporting the robustness of the conclusions with the *N*_sim_ = 10 realisations performed. The mechanism proposed above, namely that clustering acts by restricting heterotypic contact rather than by altering the mechanical properties of individual mutant cells, is tested directly in the next section.

### 3.3. Mutant spatial organisation controls heterotypic exposure and effective mechanical loading

The kinetic differences described above suggest that what sets the rate of T_2_-mediated removal is not the number of adhesion-deficient cells but the extent of their contact with the surrounding wild-type tissue. To test this directly, we quantified the heterotypic interface throughout the clearance process (figure 2). Consistent with the results of Section 3.2, randomly distributed mutants were rapidly eliminated, whereas clustered mutants persisted substantially longer over a broad range of initial mutant fractions and for both initial tissue disorders considered (*D* = 0.1 and *D* = 0.5; figure 2A,B). The total heterotypic interface *L*_het_(*t*) decreased concurrently with mutant removal (figure 2C,D). However, because *L*_het_ is intrinsically coupled to the number of surviving mutant cells, total interface length alone does not distinguish loss of interface due to clearance from changes in the geometry of the remaining mutant population. We therefore quantified the heterotypic exposure per remaining mutant, *χ*_het_ (equation (6)).

Randomly dispersed mutants showed a relatively high initial exposure to wild-type cells and were rapidly cleared (figure 2E,F). In contrast, clustered mutants started with substantially lower heterotypic exposure per cell, reflecting the shielding of cells located in the cluster interior. As clearance proceeded, *χ*_het_ progressively increased in clustered populations, consistent with the progressive erosion of the cluster and the increasing exposure of the remaining mutant cells to the surrounding wild-type tissue. Thus, the prolonged lifetime of clustered mutants is associated with their initially reduced heterotypic exposure rather than simply with their larger total interface.

Reparametrising the same trajectories by the remaining mutant fraction *φ*_mut_ (Section 3.2) reveals a clear geometrical distinction between the two spatial organisations (figure 3A,B). For a compact two-dimensional cluster, *N R*^2^ while the cluster boundary scales as *L*_het_ ∼ *R*, giving *L*_het_ *N* ^1^*^/^*^2^ and therefore *χ*_het_ ∼ *N* ^−1^*^/^*^2^. Accordingly, progressive erosion of a compact cluster predicts

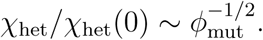

**Figure 3:**
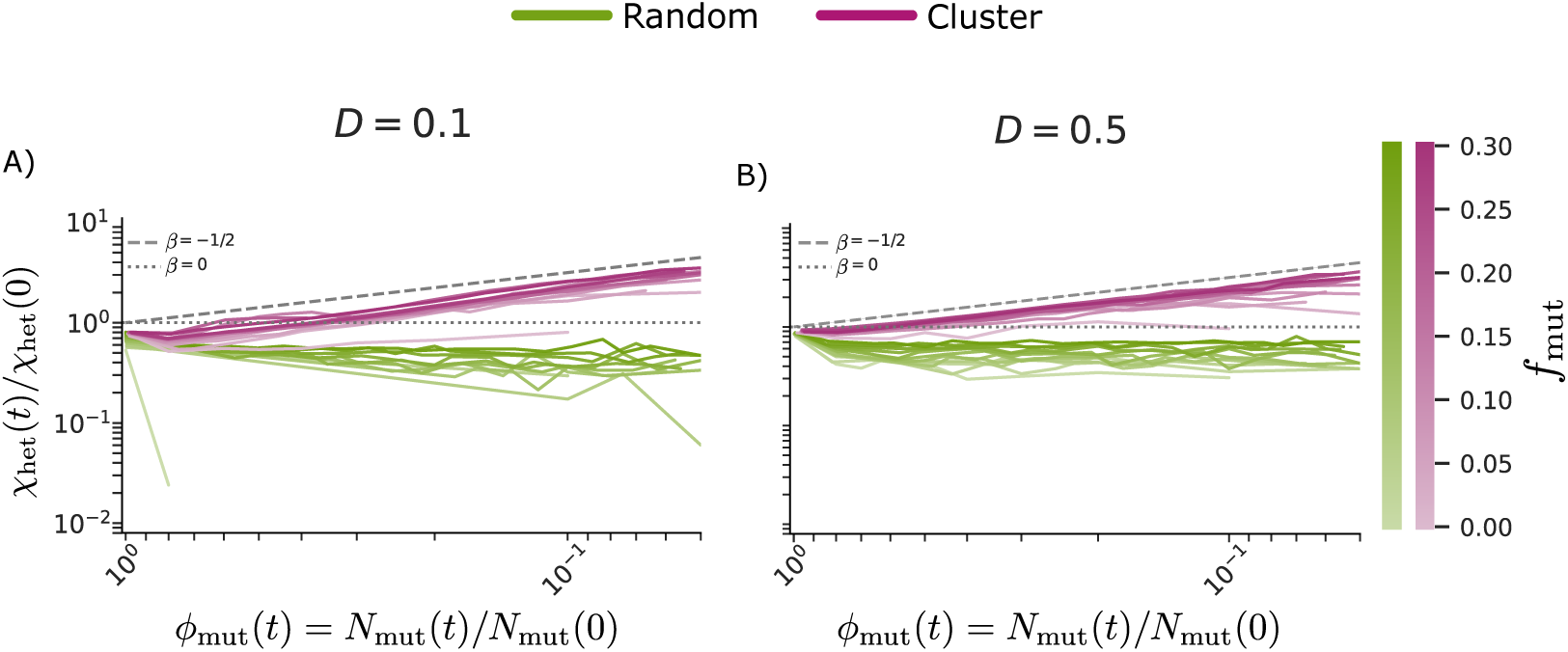
Heterotypic exposure per surviving mutant cell, rescaled by its initial value, *χ*_het_(*t*)*/χ*_het_(0), plotted against the remaining mutant fraction *φ*_mut_ = *N*_mut_(*t*)*/N*_mut_(0) on logarithmic axes, for (A) *D* = 0.1 and (B) *D* = 0.5. The horizontal axis decreases from left to right, so that trajectories are traversed in the direction of increasing clearance. Green and magenta curves correspond to randomly distributed and clustered mutant configurations, respectively; colour intensity encodes the initial mutant fraction, from *f*_mut_ = 0.025 (light) to *f*_mut_ = 0.30 (dark). The dashed line indicates the scaling *χ*_het_*/χ*_het_(0) ∼ *ϕ*_mut_*^β^* with *β* = −1*/*2, expected for the progressive erosion of a compact two-dimensional domain, and the dotted line the *β* = 0 behaviour expected for a population of approximately independent defects. After the initial transient, clustered configurations rise with an effective exponent intermediate between the two references and closer to the compact-domain value, whereas randomly distributed configurations remain close to or slightly below *β* = 0. Curves are normalised by their value at *t* = 0; the initial drop below unity reflects the contraction of *L*_het_ during the first mechanical relaxation transient, before any appreciable mutant loss. The rest of the simulation parameters are described in table 1.

The clustered trajectories show an increase in normalised exposure as *φ*_mut_ decreases, with an effective exponent that lies between the two limiting behaviours and is closer to the compact-domain value than to *β* = 0. The departure from the ideal 1*/*2 is expected, since the eroding clone is neither perfectly circular nor of uniform boundary resolution. Randomly dispersed mutants, by contrast, remain close to or slightly below the *ϕ*^0^_mut_ behaviour expected for approximately independent defects, for which heterotypic boundary length scales roughly linearly with mutant number. The erosion of a clustered population therefore follows a distinct geometrical route from the dilution of a dispersed one, and this difference accounts for the separation in clearance kinetics.

These geometric differences have a direct mechanical counterpart, but one that is not the naive expectation. Figure 4 shows the spatial distribution of the per-cell effective edge tension T̄_α_^eff^ (equation (8)), and the dynamics are illustrated in electronic supplementary material, videos S3 and S4, table 2. Because cortical contractility is identical for wild-type and mutant cells, the contractile contribution Γ(*P_α_* + *P_α_′*) is comparable across junction types at *t* = 0, and differences in T̄_α_^eff^ between them therefore reflect the line-tension contrast alone. At *t* = 0, the cells carrying the highest effective tension anywhere in the tissue are those in the interior of the mutant cluster: every one of their junctions is a homotypic mutant–mutant edge with Λ_mut_ = 0.68, whereas a randomly distributed mutant carries only heterotypic edges with Λ_hetero_ = 0.3775 and therefore an intermediate value, and wild-type junctions away from the interface carry Λ_WT_ = 0.075. Yet these interior cells are precisely the ones that survive longest. The absolute magnitude of the effective edge tension experienced by a mutant cell therefore does not predict the rate at which it is removed.

**Figure 4:**
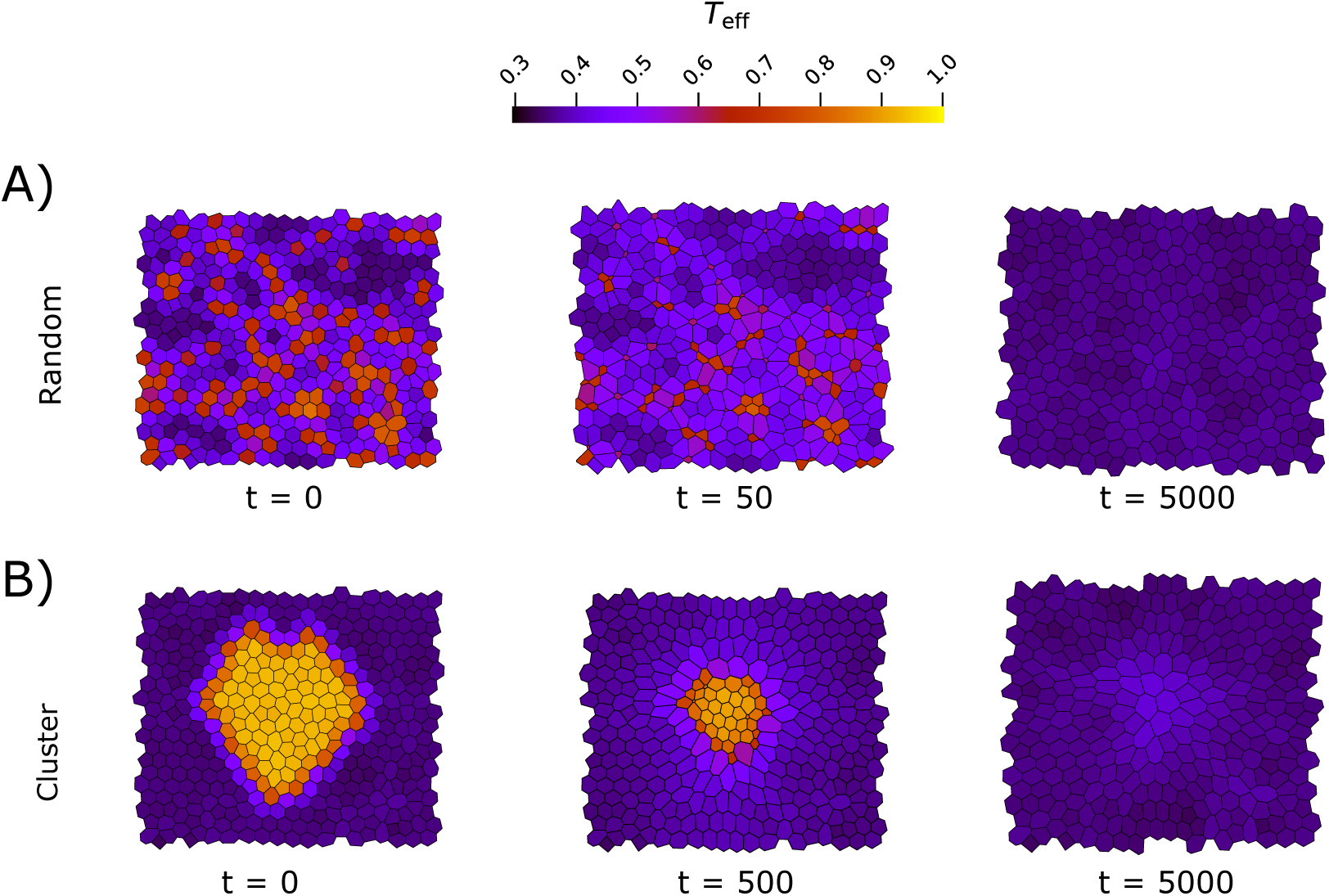
Spatial distribution of the per-cell mean effective edge tension T̄_α_^eff^ (equation (8)) during tissue evolution (*D* = 0.25, *f*_mut_ = 0.25) for randomly distributed (A, at *t* = 0, 50, and 5000) and clustered (B, at *t* = 0, 500, and 5000) mutant configurations. Warmer colours correspond to higher effective tension. At *t* = 0 the interior of the mutant cluster carries the highest effective tension in the tissue, since all of its junctions are homotypic mutant–mutant edges with Λ_mut_ *>* Λ_hetero_, whereas randomly distributed mutants, all of whose junctions are heterotypic, carry an intermediate value. Following mutant clearance, random configurations relax to an essentially uniform wild-type tension field, while clustered configurations retain a region of mildly elevated effective tension where the clone was located. The rest of the simulation parameters are described in table 1.

**Figure 5:**
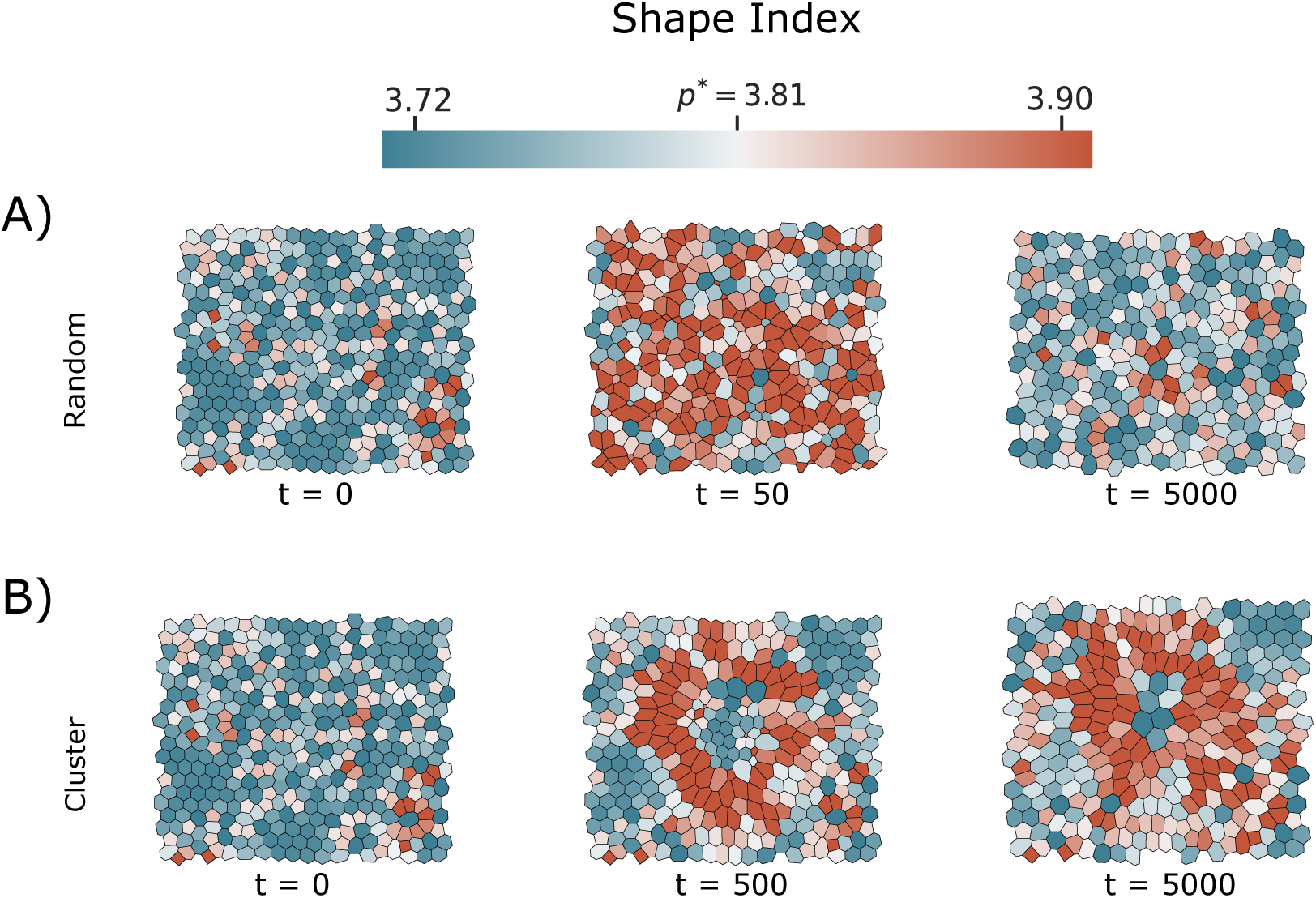
Spatial distribution of the cellular shape index *p* during tissue evolution (*D* = 0.25, *f*_mut_ = 0.25) for randomly distributed (A, at *t* = 0, 50, and 5000) and clustered (B, at *t* = 0, 500, and 5000) mutant configurations. Warmer colours correspond to higher values of *p*, indicating more elongated cells. The colour bar marks *p*^∗^ 3.81 as a reference jamming-unjamming threshold derived for homogeneous tissues; in this heterogeneous system it serves as an orientation rather than a precisely calibrated boundary. In random configurations the shape-index elevation is broadly distributed across the tissue but short-lived, having largely relaxed by *t* = 5000; in clustered configurations it is organised around the site of the clone, persists longer and extends further into the wild-type tissue. The rest of the simulation parameters are described in table 1.

What distinguishes the two configurations is instead the *contrast* in effective tension between a cell and its surroundings. In the interior of a cluster the tension field is locally uniform: a cell and all of its neighbours are mechanically equivalent, so no individual cell is preferentially compressed relative to the others, and the patch relaxes collectively to a more contracted but stable packing. Elimination through a T_2_ event requires a mechanical asymmetry between a cell and its immediate environment, which in this model arises predominantly where junctions of different line tension meet, that is, at the heterotypic interface. Randomly distributed mutants experience this asymmetry on every junction from the outset, whereas in a cluster it is confined to a boundary layer whose extent per surviving mutant cell is precisely *χ*_het_. The outside-in erosion of the cluster and the concurrent rise of *χ*_het_ thus correspond to the progressive advance of this mechanical contrast from the boundary into the interior, and it is this advance, rather than any change in the properties of individual mutant cells, that sets the clearance timescale.

An immediate consequence is that the wild-type cells bordering a clustered clone are exposed to this mechanical contrast for as long as the clone survives, which, as shown in Section 3.2, is several times longer than in random configurations. The following two sections examine what this prolonged exposure does to the geometry and the topology of the surrounding tissue.

### 3.4. Shape-index elevations at local and global scales

To characterise how shape-index elevations associated with T_2_ events extend through the epithelium, we monitored the cellular shape index, *p* = *P/ √A*, for each cell throughout the simulations. This quantity serves as a geometric proxy for unjamming-associated changes in tissue mechanical state [11, 12, 10]. Figure 5 shows an example of the spatial distribution of *p* during tissue evolution for random (panel A) and clustered (panel B) mutant configurations with *D* = 0.25 and *f_mut_* = 0.25, and the dynamics are illustrated in electronic supplementary material, videos S5 and S6, table 2. In both cases, regions of elevated shape index emerge around T_2_ sites, reflecting transient cell elongation consistent with local mechanical loading [24].

The spatial extent of this response depends on the mechanical state of the tissue. In more ordered, solid-like tissues, either because the initial packing is less noisy or because the fraction and distribution of mutant extrusions introduce only limited disorder, the tissue resists collective rearrangement [28, 10, 34]. Under these conditions, the elevated shape index remains concentrated near T_2_ sites, decaying rapidly with distance and remaining largely confined to the first ring of wild-type neighbours. By contrast, in tissues displaying geometric signatures of reduced rigidity, arising from noisier initial conditions or from a larger and/or more clustered mutant population, the epithelium can redistribute deformation more effectively through cooperative rearrangements [12]. As a result, the local shape-index elevation is less pronounced, but its signature extends over a broader region. In other words, ordered tissues localise shape-index elevation, whereas tissues closer to the unjamming-like threshold allow it to spread.

This framework also accounts for the difference between random and clustered mutant configurations. In random tissues the elevation is spatially incoherent and short-lived: because removal events are rapid and independent, they occur throughout the tissue at once and relax together, leaving no organised structure behind (figure 5A). In clustered tissues, by contrast, the elevated-*p* cells form a connected region around the eroding clone that persists as the clone is removed, promoting more persistent deviations from the preferred hexagonal geometry and extending the affected region (figure 5B). Nevertheless, this extension of shape-index elevations remains limited in tissues close to the jammed state, where elevated shape index values decay rapidly beyond the T_2_ site, consistent with the suppression of long-range rearrangements in solid-like epithelial packings [11, 10, 34].

A central question in epithelial mechanics is whether adhesion defects drive shape-index elevations across the whole tissue, or whether their mechanical impact remains confined to the immediate neighbourhood of the defects [28, 10]. To address this, we analysed the temporal evolution of the mean cellular shape index both globally *p* _global_ and locally *p* _local_. In vertex models, when *p* exceeds a critical threshold (*p*^∗^ 3.81), the tissue displays geometric signatures associated with reduced rigidity [28, 11, 29]. We note that this threshold was established for homogeneous tissues [11, 33]; in the heterogeneous setting studied here, it serves as a reference value for comparison rather than a precisely calibrated boundary.

The row *f*_mut_ = 0 in figure 6 C and D heatmaps provide the wild-type baseline, showing that *p* increases monotonically with *D* in the absence of mutant cells, consistent with [28]. This baseline allows the contribution of adhesion-deficient cells to be assessed directly relative to the disorder-driven mechanical state of a pure wild-type tissue.

**Figure 6:**
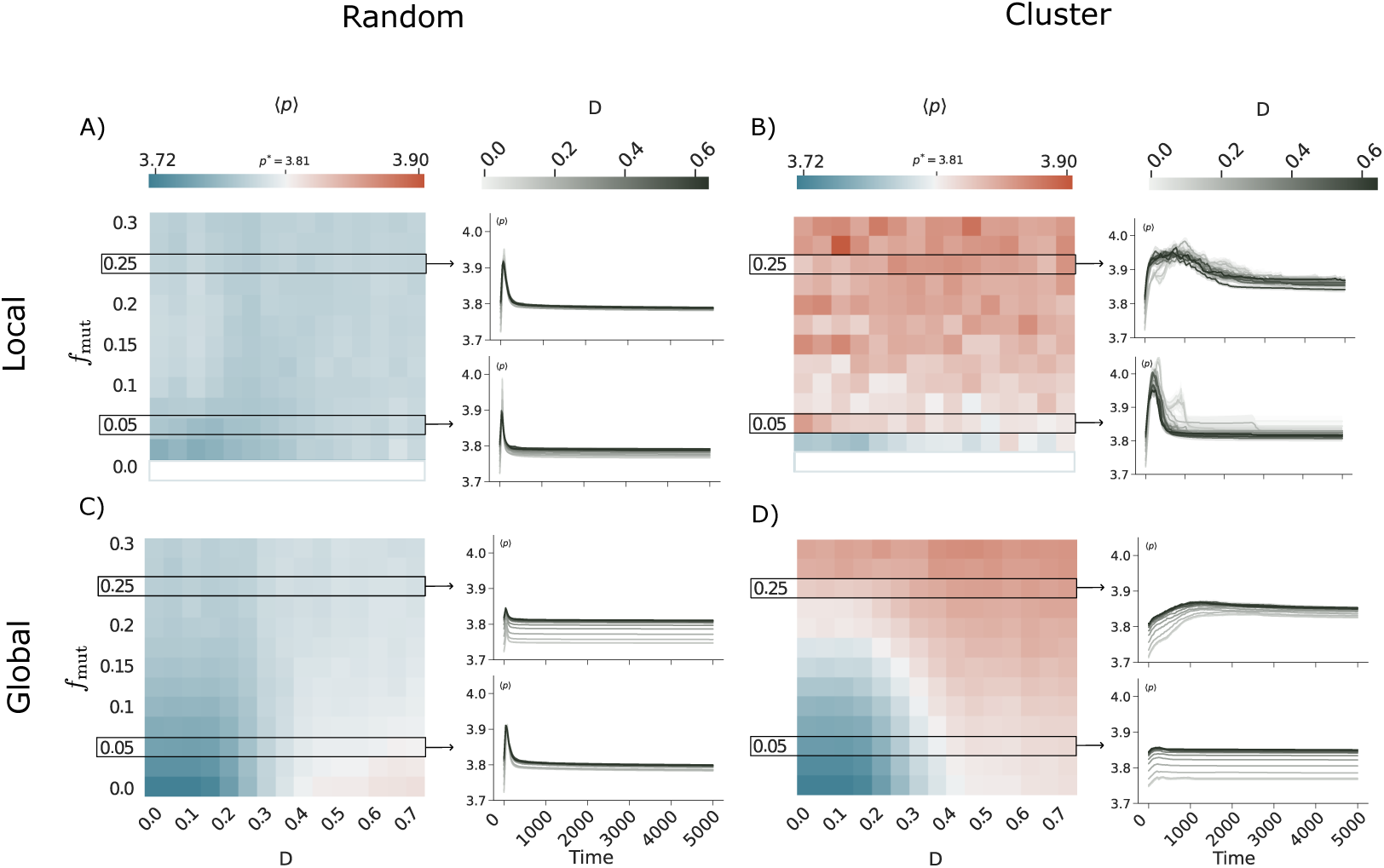
Shape-index response of the tissue to adhesion-deficient cells at two spatial scales, for random (A, C) and clustered (B, D) spatial distributions. Panels (A, B) show the local response *p* _local_, averaged over the mutant-neighbourhood cells. Panels (C, D) show the global response *p* _global_, averaged over all cells. Heatmaps show the mean shape index across the parameter space (*f*_mut_*, D*), evaluated after T_2_-mediated mutant removal at *t* = 15000 for both random and clustered configurations, exceeding *t*_5%_ for both spatial arrangements and corresponding to a mechanically equilibrated state (Supplementary Figure S1). All four panels share the same colour scale (top), where blue and red indicate shape-index values below and above *p*^∗^, respectively. Right-hand plots display the temporal evolution of *p* for representative parameter combinations (*f*_mut_ = 0.05 and 0.25, horizontal boxes in the heatmaps), with greyscale intensity encoding the initial mesh distortion *D*. Simulations were run to *t* = 15000, the time at which the heatmaps are evaluated for both configurations; the time axis of the right-hand plots is truncated at *t* = 5000 to resolve the extrusion transient. The rest of the simulation parameters are described in table 1.

To compare the post-extrusion mechanical state across configurations, the heatmaps in Figure 6 are evaluated at a common time, t = 15000, which exceeds *t*_5%_ for both spatial arrangements (Section 3.2) and, as confirmed by the total tissue energy remaining constant well before this time across the explored parameter space (Supplementary Figure S1), corresponds to a mechanically stationary state of the deterministic dynamics rather than a transient. Evaluating both configurations at the same time point avoids introducing any asymmetry between random and clustered arrangements when comparing their residual mechanical and geometric state, while accompanying time-series panels (right-hand plots) display the full temporal evolution of <*p*>, including the earlier transient peak characteristic of random configurations.

Figure 6 compares the temporal evolution of *p* for random and clustered distributions across both scales. At the global scale, random configurations generate sharp but transient peaks in ⟨*p*⟩_global_ associated with rapid T_2_ events, after which the tissue returns below *p*^∗^: at *t* = 15000 the random heatmap lies below *p*^∗^ over essentially the whole parameter space (figure 6C). Clustered mutants, by contrast, produce elevations of *p* _global_ that do not relax within the simulated window, and the clustered heatmap lies *above p*^∗^ over most of the (*f*_mut_*, D*) plane at the same time point (figure 6D). At the local scale, the contrast between configurations is even more pronounced. For random distributions (figure 6A) the shape index of the wild-type cells neighbouring the mutants, ⟨*p*⟩ _local_, spikes sharply at the moment of each T_2_ event but returns rapidly to baseline, and by *t* = 15000 it lies below *p*^∗^ throughout the parameter space, reflecting the mutually independent nature of the perturbations [24]. For clustered distributions (figure 6B), ⟨*p*⟩ _local_ remains persistently elevated throughout the T_2_ removal process and remains above *p*^∗^ over most of the parameter space at *t* = 15000, indicating that the mutant-neighbourhood cells experience sustained shape-index elevation. Notably, this local elevation appears before any rise in ⟨*p*⟩ _global_ (figure 6D), suggesting that the cluster boundary region acts as the primary site of shape-index elevation before any global signal becomes detectable; we have not measured a spatial profile of *p*, so this ordering is temporal rather than a demonstration of spatial propagation.

This relationship between scale and tissue order clarifies the coupling between the two measurements. When the tissue is well within the jammed regime, ⟨*p*⟩ _local_ captures essentially the full extent of the shape-index elevation, while ⟨*p*⟩ _global_ remains largely unaffected [28]. It is only when the tissue is driven closer to *p*^∗^ that shape-index elevations extend beyond the first-neighbour shell and produce a measurable global response [12, 10]. The divergence between the two signals therefore acts as a geometric indicator of proximity to the unjamming-like threshold: a large ⟨*p*⟩ _local_ with a small ⟨*p*⟩ _global_ indicates a tissue that is locally perturbed but globally near its homeostatic state, whereas convergence of the two signals suggests the onset of tissue-wide shape-index elevation consistent with unjamming-like behaviour. This threshold behaviour is controlled by two parameters that act through distinct mechanisms. Initial disorder, *D*, reduces the energetic cost of topological rearrangements globally by pre-distorting the lattice away from its optimal hexagonal geometry [11, 10, 34]. Mutant fraction, *f*_mut_, by contrast, introduces localised sources of mechanical stress whose cumulative effect grows with their number [33]. The spatial organisation of mutant cells then determines whether these sources act independently, by producing additive but uncorrelated perturbations that remain local; or cooperatively, by generating a coherent pattern of shape-index elevation that extends beyond the first-neighbour shell even at moderate *f*_mut_. In this sense, increasing *f*_mut_ and increasing *D* produce qualitatively similar effects on the global shape index, but clustering acts as a spatial amplifier that produces tissue-wide shape-index elevations consistent with unjamming-like signatures at lower values of both parameters than either alone would require.

These results establish that spatial organisation is an additional structural variable that modulates the mechanical response of the tissue independently of mutant density in our simulations: while mutant fraction determines the overall magnitude of the perturbation, the spatial arrangement of those cells governs both its duration and the extent to which shape-index elevations spread beyond the immediate T_2_ sites.

As an additional, dynamical corroboration of this geometric picture, we computed the cumulative mean-squared displacement of the mutant neighbourhood wild-type cells, MSD_0_(*t*), as defined in Supplementary Section S1.1 (Supplementary Figure S3). Clustered configurations show cumulative displacements roughly an order of magnitude larger than random configurations at matched mutant fraction and initial disorder, with the difference most pronounced at high mutant fraction. In both cases MSD_0_(*t*) saturates at a finite plateau rather than growing indefinitely, indicating a bounded structural rearrangement of the surrounding tissue rather than sustained fluid-like motion; the prolonged shape-index elevation in clustered configurations therefore reflects a genuine, if finite, displacement of the wild-type tissue rather than a purely static geometric signature.

### 3.5. Spatial organisation induces persistent topological disorder

Beyond deforming cell shapes, the extrusion of adhesion-deficient cells also disrupts the connectivity of the epithelial network [24, 57]. Figure 7 shows the spatial distribution of the number of neighbours for each cell at successive time points, for random (panel A) and clustered (panel B) configurations with *D* = 0.25 and *f_mut_* = 0.25, and the dynamics are illustrated in electronic supplementary material, videos S7 and S8, table 2. Under homeostatic conditions, the majority of cells are hexagonal (*n* = 6); deviations indicate topological rearrangements induced by extrusion events [1, 34].

**Figure 7:**
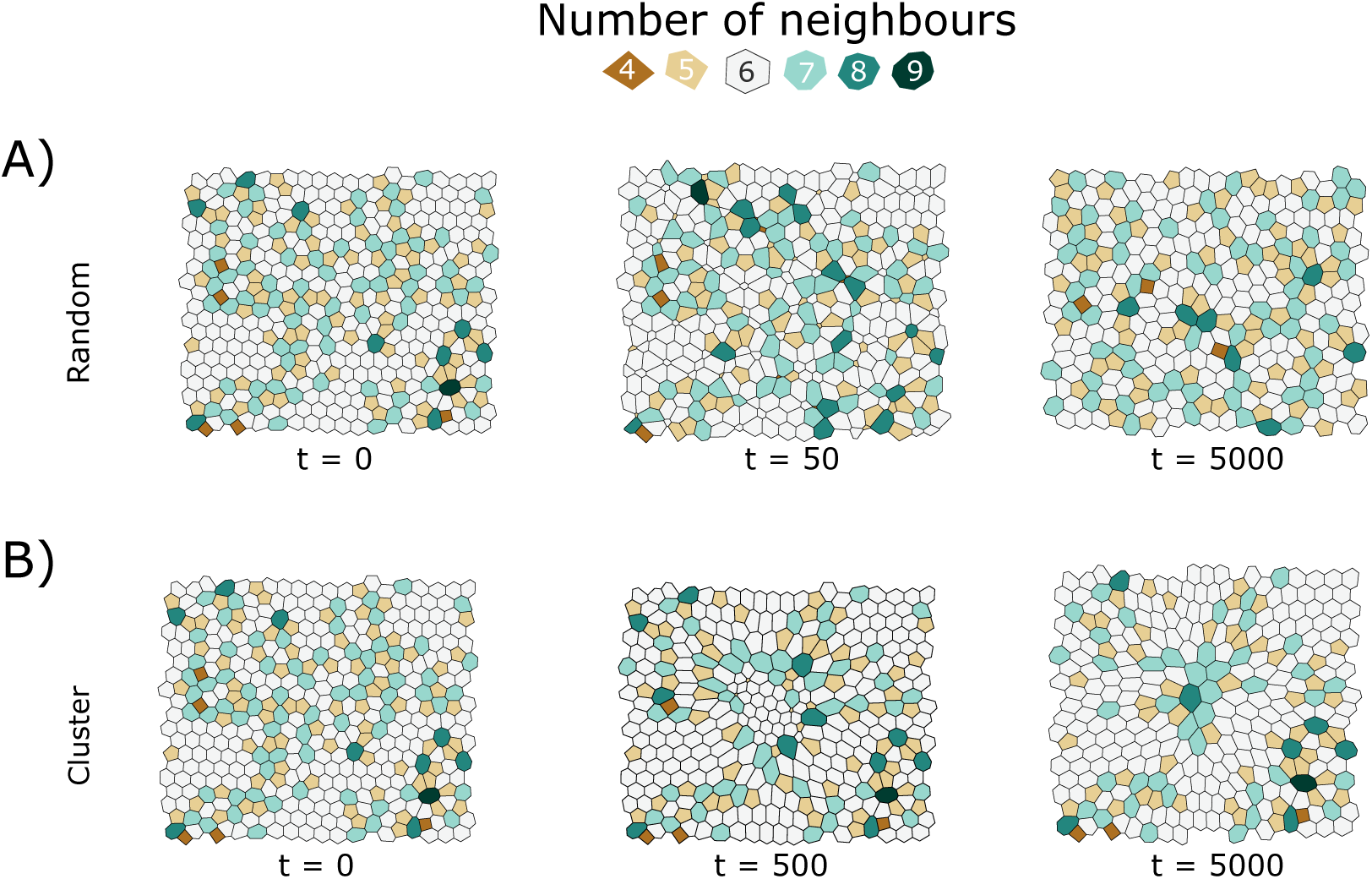
Spatial distribution of the number of neighbours per cell during tissue evolution (*D* = 0.25, *f*_mut_ = 0.25) for randomly distributed (A, at *t* = 0, 50, and 5000) and clustered (B, at *t* = 0, 500, and 5000) mutant configurations. Colours indicate the number of sides of each polygon as shown in the legend. Hexagonal cells (*n* = 6) dominate the homeostatic state; non-hexagonal cells mark regions of topological rearrangement driven by extrusion events. In random configurations the non-hexagonal cells remain scattered throughout the tissue; in clustered configurations they form a contiguous, persistent zone at the site of the clone, including cells of high polygon class where several neighbours converge on a common vertex. The rest of the simulation parameters are described in table 1.

In random configurations, cells with anomalous neighbour numbers remain spatially scattered and do not organise into any larger structure, as each T_2_ event affects only its immediate neighbourhood before the tissue relaxes [34]. In clustered configurations, the coordinated T_2_-mediated removal of adjacent mutant cells produces a spatially extended and persistent zone of topological disorder around the cluster boundary, characterised by a broad shift of the local polygon distribution towards classes above six. This topological imprint persists within the simulated time window after the last mutant cell has been removed, suggesting that clustered configurations leave a structural imprint on the mutant-neighbourhood wild-type cells that random configurations do not (Supplementary Figure S4). The temporal evolution of polygon class frequencies quantifying this effect is shown in figure 8.

**Figure 8:**
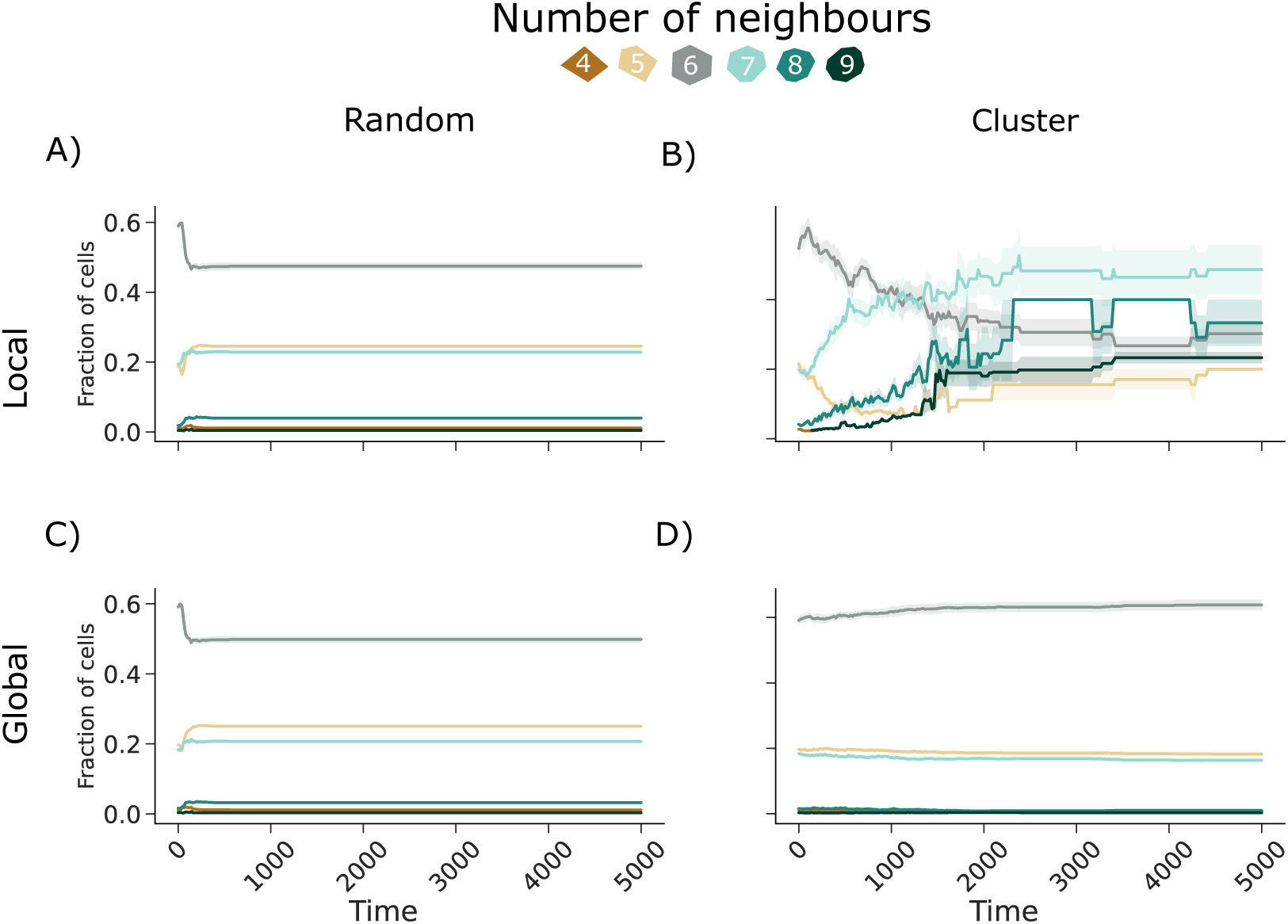
Temporal evolution of polygon class frequencies (fraction of cells with *n* neighbours) for randomly distributed (A, C) and clustered (B, D) mutant configurations (*D* = 0.25, *f*_mut_ = 0.25). Panels (A, B) show the local response (mutant-neighbourhood wild-type cells); panels (C, D) show the global response (all cells). Colours indicate polygon class as shown in the legend. Curves represent the mean over *N*_sim_ = 10 independent simulations; shaded bands indicate the standard error of the mean, which is appreciably wider for the clustered local set (B) because it is built up progressively as the clone is eroded. Simulations were run to *t* = 15000; the time axis is truncated at *t* = 5000 to resolve the dynamics. The rest of the simulation parameters are described in table 1.

To quantify these qualitative observations systematically across the full parameter space, we computed the mean absolute deviation from hexagonal packing, ⟨Δ_hex_⟩, defined as the mean over all cells of *n_i_* 6 where *n_i_* is the number of neighbours of cell *i*. In epithelial monolayers near mechanical equilibrium, cells tend towards hexagonal packing due to geometric constraints and force balance [1, 54, 34]; ⟨Δ_hex_⟩ thus measures the degree to which T_2_-mediated removal displaces the tissue from this reference state across the parameter space defined by *f*_mut_ and *D*.

Figure 8 quantifies the temporal evolution of polygon class frequencies for both configurations. In random configurations the shift away from hexagonal packing is modest and closely similar at the two scales, and the resulting distribution is approximately symmetric about *n* = 6; it is established early and then persists unchanged for the rest of the window. In clustered configurations, by contrast, the local distribution shifts asymmetrically towards classes above six, with heptagons becoming the most abundant class and with octagons and nonagons acquiring substantial populations that have no counterpart in random configurations, indicative of a persistent loss of trans-lational order [57, 24]. This asymmetric signature is consistent with the persistent shape-index elevation observed in *p* _local_, and is essentially absent from the corresponding global distributions. The polygon distributions at *t* = 0 and at the end of the simulation for representative parameter combinations are shown in Supplementary Figure S4.

Figure 9 shows the resulting heatmaps for random (panels A, C) and clustered (panels B, D) configurations. Here, local measurements (panels A–B) refer to the mutant-neighbourhood wild-type cells (wild-type neighbours of mutant cells, updated while each mutant cell is present and fixed once it is cleared), and global measurements (panels C–D) refer to all cells in the tissue, consistent with the local/global distinction defined in Section 2.4. Time evolution plots show that, once mutant removal is complete in both configurations, a striking asymmetry emerges: locally, clustered configurations leave behind a substantially more disordered topological landscape than random distributions for equivalent parameter values.

**Figure 9:**
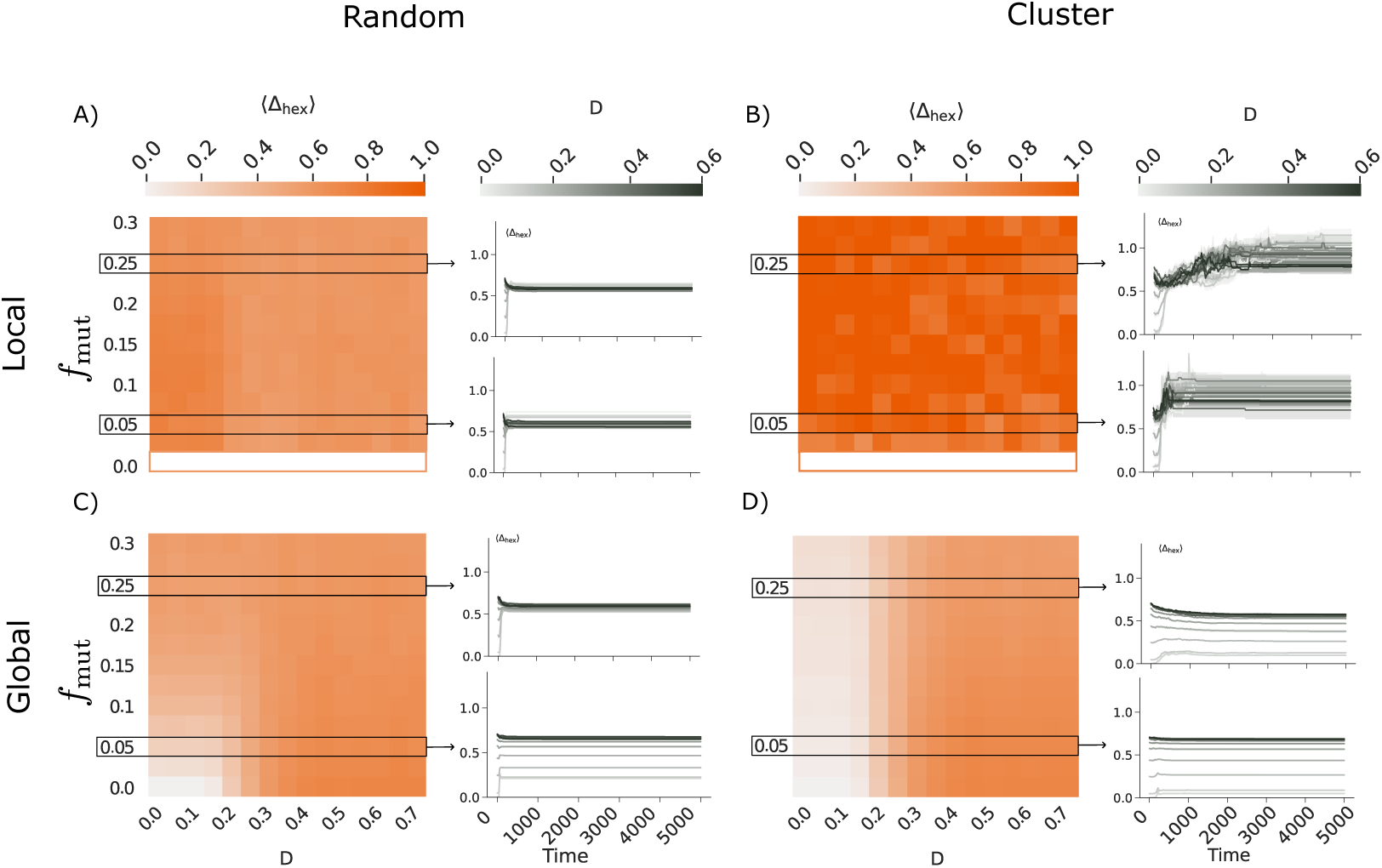
Heatmaps of the mean absolute deviation from hexagonal packing, ⟨Δ_hex_⟩ = ⟨|*n_i_-*6|⟩, across the parameter space (*f*_mut_*, D*) for randomly distributed (A, C) and clustered (B, D) mutant configurations, evaluated after T_2_-mediated mutant removal at *t* = 15000 for both random and clustered configurations, exceeding *t*_5%_ for both spatial arrangements and corresponding to a mechanically equilibrated state (Supplementary Figure S1). Darker colours indicate greater deviation from hexagonal packing. Local measurements (A, B) are restricted to the mutant-neighbourhood wild-type cells (wild-type neighbours of mutant cells, updated while each mutant cell is present and fixed once it is cleared); global measurements (C, D) are averaged over all cells. Right-hand plots show the temporal evolution of ⟨Δ_hex_⟩ for representative parameter combinations (*f*_mut_ = 0.05 and 0.25, horizontal boxes in the heatmaps), with greyscale intensity encoding the initial mesh distortion *D*. Simulations were run to *t* = 15000, the time at which the heatmaps are evaluated for both configurations; the time axis of the right-hand plots is truncated at *t* = 5000 to resolve the extrusion transient. At the local scale, clustered configurations produce substantially greater topological disorder than random distributions. At the global scale the ordering is reversed, because random removal events are distributed throughout the tissue whereas clustered removal is confined to the neighbourhood of the clone. The rest of the simulation parameters are described in table 1.

### 3.6. Mutant clearance leaves a residual mechanical loading of the surrounding tissue

The exposure mechanism of Section 3.3 accounts for the clearance kinetics of the mutant population, but not for why the elevated shape index (Section 3.4) and the topological disorder (Section 3.5) persist in the surrounding wild-type tissue after the mutant cells themselves are gone. To address this, we followed the effective edge tension at the local and global scales defined in Section 2.5 (figure 10). Note that the global average is taken over all edges in the tissue, so that while mutant cells survive it also includes their homotypic and heterotypic junctions; the local average is restricted throughout to the edges of the mutant-neighbourhood wild-type cells.

**Figure 10:**
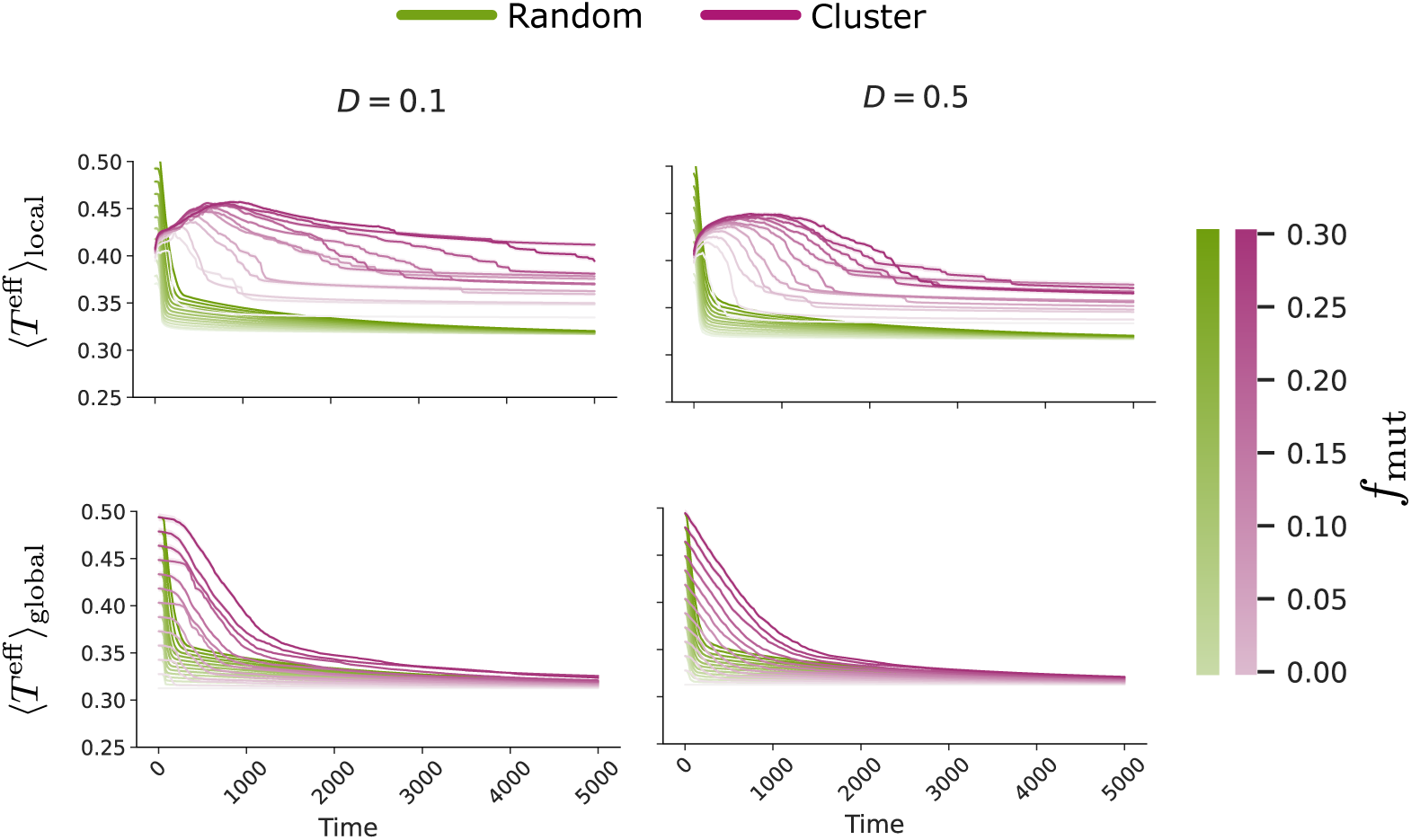
Temporal evolution of the mean effective edge tension at the local (top row) and global (bottom row) scales, for low (*D* = 0.1, left) and high (*D* = 0.5, right) initial mesh distortion. Local values are averaged over the edges of the mutant-neighbourhood wild-type cells; global values over all edges in the tissue (Section 2.5). Green and magenta curves correspond to randomly distributed and clustered mutant configurations, respectively. Colour intensity encodes the initial mutant fraction, from *f*_mut_ = 0 (light) to *f*_mut_ = 0.30 (dark); the local panels contain no *f*_mut_ = 0 curve, since the mutant-neighbourhood set is undefined in the absence of mutant cells. At the global scale both configurations relax to a common wild-type baseline, whereas at the local scale clustered configurations remain systematically elevated above random configurations throughout the interval shown, and are still decaying slowly at *t* = 5000. Each curve represents the mean over *N*_sim_ = 10 independent simulations; shaded bands indicate the standard error of the mean (SEM). Simulations were run to *t* = 15000; the time axis is truncated at *t* = 5000 to resolve the dynamics. The rest of the simulation parameters are described in table 1.

At the global scale the two configurations behave alike: ⟨*T* ^eff^⟩ _global_ decays from its initial value and converges onto a common wild-type baseline, essentially independently of spatial arrangement and mutant fraction, consistent with the relaxation of the total tissue energy reported in Supplementary Figure S1. It is the local signal that separates the two configurations. In random configurations, ⟨*T* ^eff^⟩ _local_ falls rapidly and settles onto the same baseline as the global signal, consistent with the transient, spatially confined shape-index and topological perturbations reported in Sections 3.4 and 3.5. In clustered configurations, by contrast, ⟨*T* ^eff^⟩ _local_ first rises during the early stage of cluster erosion and then decays only slowly, remaining systematically above both the random curves and the global baseline throughout the interval shown, with the separation increasing with *f*_mut_. At *D* = 0.1 and the highest mutant fraction, for example, the local signal peaks at ≃0.50 near *t*≃900 and has fallen only to≃0.46 by *t* = 5000, against a global baseline of≃0.33; the residual loading therefore decays on a timescale much longer than that of mutant clearance itself, which is complete by *t* ≃2000 (Supplementary Figure S2).

The wild-type cells that surrounded the clone are therefore left carrying a residual mechanical load that outlasts the clone itself. This provides a mechanical correlate of the persistent shape-index elevation and of the excess of high-order polygon classes documented in the preceding sections, rather than a separate phenomenon: the same cells that remain elongated and topologically irregular are the ones whose junctions remain under elevated effective tension.

Taken together, figures 2, 3 and 10 indicate that the exposure imbalance between clustered and randomly distributed adhesion-deficient cells accounts for the delayed T_2_-mediated clearance of clustered mutants (Section 3.2), the persistent local elevation of the shape index (Section 3.4), and the sustained topological disorder observed after mutant clearance (Section 3.5). We therefore interpret these three signatures not as independent evidence for a tissue-wide rigidity transition, but as correlated geometric and mechanical consequences of a single underlying mechanism.

## 4. Discussion and Conclusions

In this study, we have employed a two-dimensional vertex model to show that the spatial organisation of adhesion-deficient cells modulates the geometric signatures of epithelial unjamming, governing mutant clearance kinetics, the extent of shape-index elevation, and the persistence of topological disorder. Previous studies have shown that cell-level mechanical heterogeneity can alter rigidity-transition behaviour [39, 40]. Here, rather than addressing the effect of mechanical heterogeneity itself, we focus on how the spatial arrangement of adhesion-deficient cells shapes the tissue response. Our results establish that the spatial *arrangement* of those cells modulates three equally important dimensions of the mechanical response: its duration, the extent to which shape-index elevations spread beyond the immediate T_2_ sites, and the persistent topological disorder left in the mutant-neighbourhood tissue.

### Heterotypic interface exposure sets the clearance timescale

The differences between random and clustered configurations reported here follow from a single geometric quantity: the heterotypic interface exposure per surviving mutant cell, *χ*_het_ = *L*_het_*/N*_mut_. In a compact clone *N* ∼ *R*^2^ while the heterotypic boundary scales as *L*_het_ ∼ *R*, so *χ*_het_ is initially small and grows as the clone is eroded, with a measured exponent between *ϕ*_mut_^-1/2^ and *ϕ*_mut_^0^ and closer to the former, whereas in a dispersed population it remains close to its initial value (Section 3.3). Because T_2_-mediated removal in this model is driven by the mechanical asymmetry between a cell and its immediate neighbours, and in this model such asymmetry is by far the largest where junctions of different line tension meet, *χ*_het_ is the natural control variable for the clearance rate.

It is worth emphasising that this is a statement about mechanical contrast rather than magnitude. Cells in the interior of a clone carry the highest absolute effective edge tension anywhere in the tissue, since all of their junctions are homotypic Λ_mut_ edges, and yet they are the last to be removed; an interpretation based on the absolute interfacial tension of mutant cells would therefore predict the opposite ordering to the one we observe. In this respect our results connect directly to work on differential interfacial tension and boundary maintenance between mechanically distinct populations [6, 51, 52, 53], where it is likewise the tension differential at heterotypic contacts, rather than the tension of either population in isolation, that governs the outcome. A direct corollary is that the wild-type tissue bordering a clone remains mechanically loaded for as long as the clone survives: the local effective edge tension of the mutant-neighbourhood wild-type cells stays elevated above the global baseline well after the last mutant cell has been removed (figure 10), providing the mechanical counterpart of the geometric and topological signatures discussed next.

### Relation to generic inclusion relaxation

Framed in this way, the clustered configuration is closely related to the relaxation of a compact inclusion embedded in a surrounding medium, a situation with qualitative analogies to defect closure and wound-healing mechanics [3], although here the inclusion is removed cell by cell through T_2_ events within a confluent monolayer rather than by closure of an open gap. We regard this correspondence as a feature of the model rather than a limitation of it. In our formulation the only element specific to adhesion loss is the parametrisation of Λ*_ij_* in equations (2)–(3); no additional adhesion-specific dynamical rule is imposed. The interface-length argument developed above is correspondingly generic: the *ϕ*_mut_^-1/2^ scaling *expected* for *χ*_het_ follows from the geometry of a compact two-dimensional domain and would hold for any inclusion whose boundary carries excess tension. What is specific to adhesion-deficient clones is not the mechanism but its parametrisation, that is, the magnitude and sign of the line-tension contrast generated by E-cadherin loss, and the fact that the inclusion is removed by the same T_2_ machinery that maintains homeostatic turnover in the surrounding tissue. Our results therefore show how a generic differential-tension inclusion-relaxation mechanism, when parametrised by biologically motivated E-cadherin-loss line tensions, produces clonal-expansion-relevant kinetic and topological signatures at the tissue scale.

### Clustering prolongs shape-index elevation

A central finding is the fundamental asymmetry between random and clustered mutant distributions in their effect on tissue shape index, figure 6. Randomly distributed adhesion defects generate localised, transient shape-index perturbations that are rapidly resolved through T_2_-mediated cell removal, allowing the tissue to return towards its homeostatic packing geometry [1, 34]. Clustered mutants, by contrast, mimic the clonal expansion observed in early-stage gastric cancer and other E-cadherin-related pathologies [15, 23], and sustain shape-index values close to *p*^∗^ for extended periods, displaying geometric signatures consistent with proximity to the unjamming threshold. This follows directly from the exposure mechanism above: the reduced heterotypic exposure of interior cluster cells delays their T_2_-mediated removal, and the surrounding wild-type tissue is therefore held under mechanical loading for correspondingly longer. This reframes the role of clonal expansion: beyond simply increasing the number of defective cells, spatial aggregation acts as a structural amplifier that sustains unjamming-like shape-index signatures at lower mutant fractions than random distributions would require [33].

### Local and global scales reveal distinct signatures

A key contribution of this work is the explicit separation of shape-index signals at local and global scales, where local refers to the mutant-neighbourhood wild-type cells (wild-type neighbours of mutant cells, updated while each mutant cell is present and fixed once it is cleared). Our analysis shows that these cells experience persistent shape-index elevation throughout the T_2_ removal process, and that this local signal appears before any rise in global *p* (figure 6, panels B and D). This ordering is consistent with a scenario in which shape-index elevations are initially more pronounced at the cluster boundary, where heterotypic interface density is highest, and subsequently propagate into the broader tissue through cascades of T_2_ events [2, 11]. In random distributions, this extension is effectively quenched because isolated T_2_ events lack the spatial coherence needed to sustain a tissue-wide shape-index perturbation [24]. The local scale therefore acts as an early geometric indicator of mechanical instability that the global shape index alone would miss [10], with potential implications for experimental detection of pre-invasive mechanical states [20].

### T_2_ events leave a persistent topological imprint

Our analysis of polygon distributions reveals that T_2_-mediated cell removal is not a perfectly restorative process [57]. In random configurations, the polygon distribution also shifts away from its initial near-hexagonal state and does not return to it within the simulated window (figure 8A,C and Supplementary Figure S4); the shift is, however, modest, approximately symmetric about *n* = 6, and of essentially the same magnitude at the local and global scales, as expected when removal events are mutually independent and distributed throughout the tissue [34]. In clustered configurations the coordinated removal of adjacent cells leaves behind a substantially more disordered local polygon distribution (Supplementary Figure S4), shifted asymmetrically towards classes above six and with no counterpart in the corresponding global distribution, which persists throughout the simulated time window even after full mutant clearance has been achieved. This persistent topological imprint, measured as deviation from hexagonal packing and excess polygon class frequencies, suggests that clonal expansion may alter the architectural organisation of the tissue in a long-lived manner [23, 13, 24]. Whether this constitutes a truly irreversible change would require simulations over much longer time scales. A tissue carrying such persistent disorder may be more susceptible to subsequent mechanical perturbations, potentially lowering the effective threshold for future shape-index elevations, though confirming this prediction would require dedicated simulations with repeated mechanical challenges.

An important caveat concerns the nature of this final state. Our dynamics are deterministic overdamped relaxations with prescribed T_1_ and T_2_ thresholds, and the regular hexagonal packing is the low-energy reference configuration of the uniform wild-type vertex model. We have verified that the disordered states we report are stationary rather than transient: the total tissue energy and its relaxation rate are flat well before the evaluation time (Supplementary Figure S1), so the residual disorder is not an artefact of measuring the tissue mid-relaxation. Stationarity under deterministic dynamics is not, however, the same as robustness: weak active fluctuations, junctional noise, or a much longer post-clearance relaxation could in principle anneal the tissue back towards a more hexagonal configuration, and we have not tested this. The disordered configurations reported here should therefore be understood as long-lived states of the deterministic protocol, whose stability under fluctuations remains to be established.

### Biological implications and outlook

Taken together, these findings (delayed clearance, prolonged shape-index elevation, scale-dependent geometric signatures, and a persistent topological imprint with its residual mechanical counterpart) converge on a unified picture in which all four are consequences of the same exposure imbalance: the geometry of clonal expansion is itself a structural determinant of the tissue’s susceptibility to mechanical destabilisation, in the sense of vertex-model shape-index criteria, independent of the biochemical identity of the defective cells. In the context of cancer progression, our model suggests that early detection strategies should consider not only the fraction of E-cadherin-deficient cells but also whether those cells form spatially correlated clusters [23, 13, 15]. A tissue with a small clustered clone may display more pronounced unjamming-like shape-index signatures than one with a larger but randomly distributed mutant population, because spatial aggregation amplifies and prolongs collective shape-index shifts [28, 10, 33].

It is worth stating explicitly how far these results reach. Three levels of conclusion can be distinguished. First, that clustering delays T_2_-mediated mutant clearance: this is robustly supported, holds throughout the parameter space explored (Section 3.2, Supplementary Figure S2), and now has an explicit geometric and mechanical explanation (Section 3.3). Second, that clustering produces a long-lived structural and mechanical disturbance of the surrounding tissue: this is supported within our simulation protocol by the persistent local shape-index elevation, the shift of the local polygon distribution towards classes above six, the residual local effective edge tension, and the order-of-magnitude larger cumulative displacement of the neighbouring wild-type cells. The shape-index, topological and displacement signatures are evaluated in configurations whose total energy has ceased to relax (Supplementary Figure S1); the local effective tension is still decaying slowly at the end of the interval we display, so its persistence is established over that interval rather than as a stationary end state. Third, that clustering produces a robust mechanical or topological memory that would survive perturbation: this we do not claim. Establishing it would require the post-clearance fluctuation or noise tests described above. Our conclusions are therefore intended at the second level, and the geometric signatures we report should be read as evidence of a prolonged, protocol-stable perturbation rather than of a durable tissue-scale memory.

Future experimental work should aim to measure shape index fields and polygon distributions at the boundary of expanding clones in controlled *in vitro* systems, where the spatial organisation of E-cadherin knockdown cells can be directly manipulated [19, 22, 20]. From a theoretical perspective, extending this framework to incorporate active motility [11], cell proliferation [58], and three-dimensional geometry [33] would allow direct comparison with *in vivo* observations of early tumour invasion.

This work provides a physical framework linking local E-cadherin deficiency to tissue-scale shape-index changes, and identifies spatial organisation as an additional structural control variable of the geometric signatures of epithelial unjamming, alongside cell density and adhesion strength, acting through the heterotypic interface exposure of the adhesion-deficient population.

### Data availability and Methods

The videos supporting this article have been uploaded as electronic supplementary material. table 2 summarises the content of each video. All simulation code used for computational modelling and data analysis is publicly available in Zenodo [59]: https://doi.org/10.5281/zenodo.22961598.

## Acknowledgments

N.B-P, V.M. and P.G. acknowledge Ministerio de Ciencia, Innovación y Universidades/European Regional Development Fund (Spain/EU) grant PGC2018-098186-B-I00 (BASIC). V.M. wants to acknowledge that the project that gave rise to these results received the support of a fellowship from Fun-dación Ramón Areces.

