## Supplemental methods and Supplemental figures 1-4 for "Spatial clustering of adhesion-deficient cells delays clearance and prolongs geometric unjamming signatures in epithelial tissues"

### S1 Additional methods

#### S1.1 Mean-squared displacement

To complement the geometric characterisation provided by the shape index with a dynamical measure of tissue rearrangement, we calculated the cumulative mean-squared displacement (MSD) of the mutant-neighbourhood wild-type cells relative to the initial tissue configuration. Denoting by $\mathcal{N}$ the set of mutant-neighbourhood wild-type cells (defined as in Section 2.4 of the main text), the position of each cell $i \in \mathcal{N}$ was defined by the centroid of its polygon, $\mathbf{r}_i(t)$, and the MSD was computed as

$$\mathrm{MSD}_0(t) = \left\langle |\mathbf{r}_i(t) - \mathbf{r}_i(0)|^2 \right\rangle_{i \in \mathcal{N}},$$

where the average is taken over the cells of $\mathcal{N}$ present at both the initial and corresponding evaluation times. This measure quantifies the cumulative displacement of the surrounding wild-type tissue from its initial configuration and is evaluated independently for each simulation and subsequently averaged across the $N_{\mathrm{sim}} = 10$ independent realisations.

### S2 Additional figures

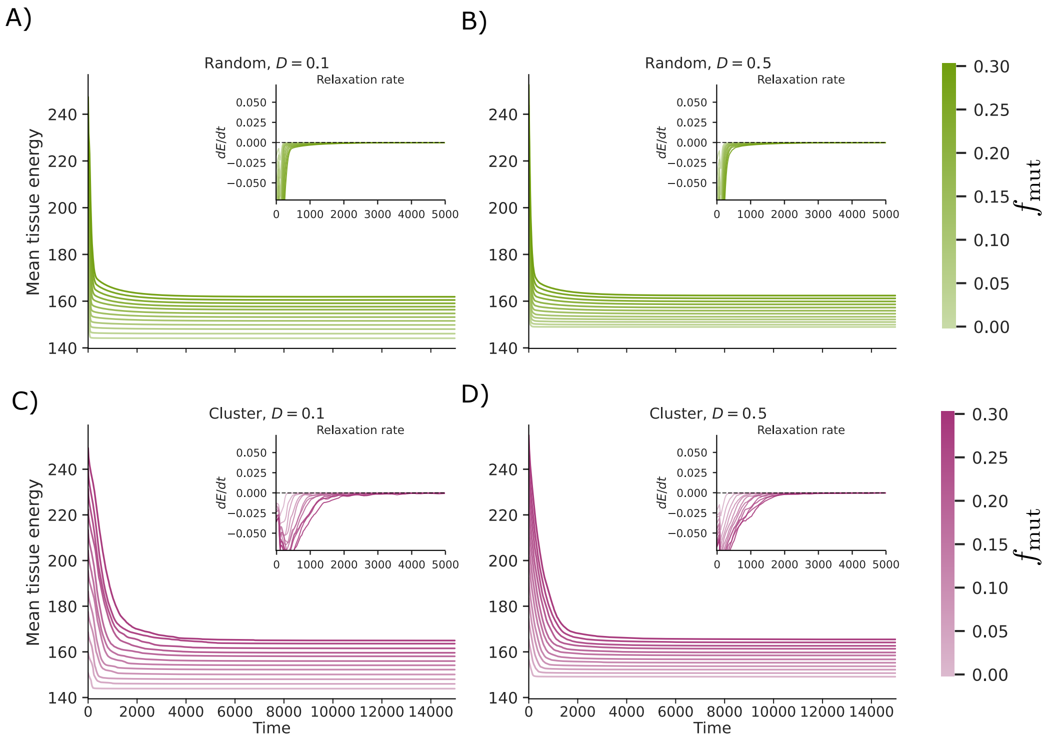


Figure S1: Mean tissue energy as a function of time for randomly distributed (A, B) and clustered (C, D) mutant configurations, at low ($D = 0.1$; A, C) and high ($D = 0.5$; B, D) initial mesh distortion. Colour intensity encodes the mutant fraction $f_{\mathrm{mut}}$; insets show the relaxation rate $dE/dt$. In all cases the energy relaxes to a stationary plateau, with $dE/dt$ indistinguishable from zero on the scale of the inset by $t \approx 2000$ and well before the evaluation time $t = 15000$ used in the main text. The rest of the simulation parameters are described in table 1 of the main text.

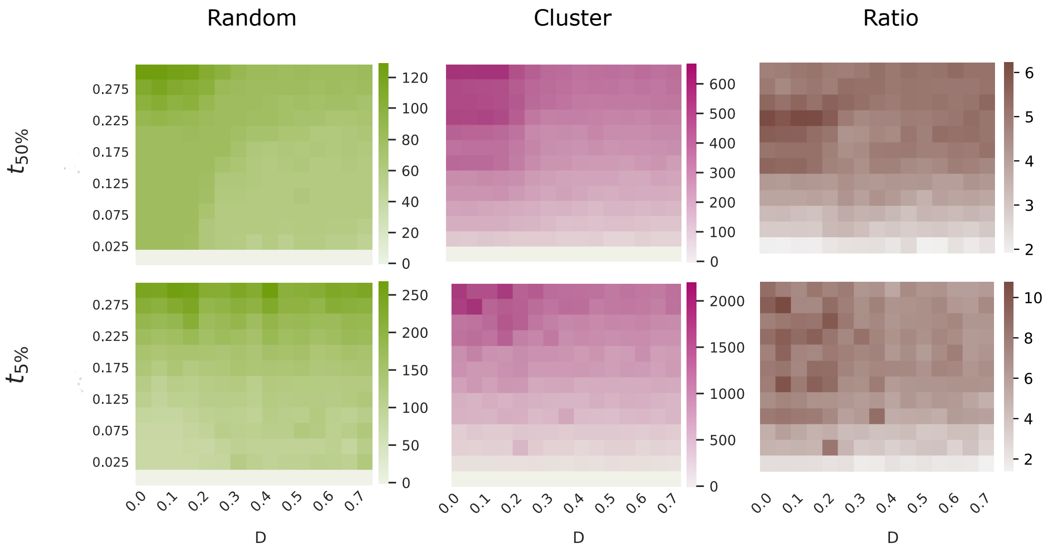


Figure S2: Time at which 50% ($t_{50\%}$, top row) and 5% ($t_{5\%}$, bottom row) of adhesion-deficient cells remain in the tissue, as a function of initial tissue distortion $D$ and mutant fraction $f_{\mathrm{mut}}$, for randomly distributed (left column) and clustered (middle column) configurations; the right column shows the ratio $t^{\mathrm{cluster}}/t^{\mathrm{random}}$. The ratio exceeds 1 throughout the parameter space and exceeds 2 for $t_{50\%}$. Clearance is complete well before the evaluation time used in the main text: $t_{5\%}$ does not exceed $\simeq 2000$ in any configuration, so that all post-extrusion observables are evaluated at $t = 15000$, at least sevenfold later than the slowest clearance we observe. The rest of the simulation parameters are described in table 1 of the main text.

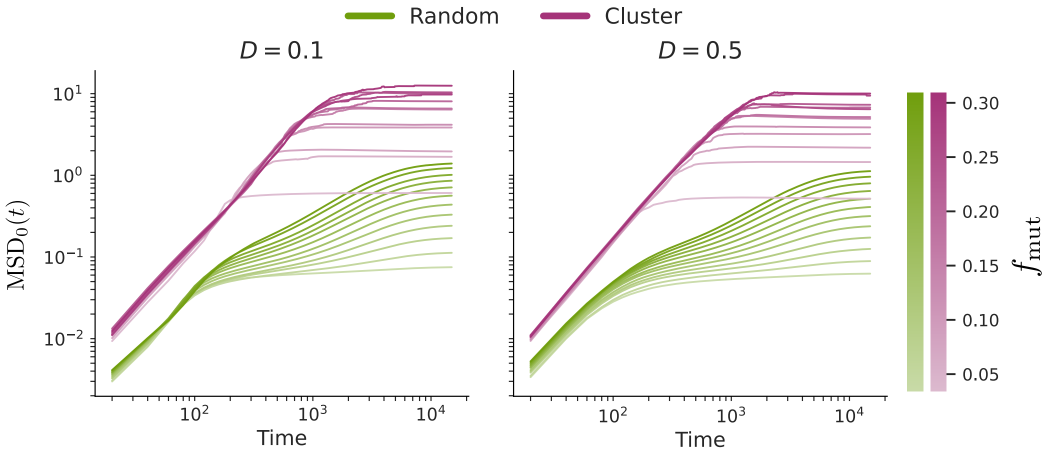


Figure S3: Cumulative mean-squared displacement of the mutant-neighbourhood wild-type cells, $\mathrm{MSD}_0(t)$ (defined in Section S1.1), for randomly distributed (green) and clustered (magenta) mutant configurations, at low ($D = 0.1$, left) and high ($D = 0.5$, right) initial mesh distortion, on logarithmic axes. Colour intensity encodes the mutant fraction $f_{\mathrm{mut}}$. In both panels, clustered configurations display cumulative displacements roughly an order of magnitude larger than random configurations at matched mutant fraction and initial disorder. The ratio between the two is comparable across $f_{\mathrm{mut}}$; it is the absolute separation that grows with mutant fraction. In all cases $\mathrm{MSD}_0(t)$ saturates at a finite plateau, indicating a bounded rearrangement of the surrounding tissue, and providing a dynamical corroboration of the shape-index and topological signatures reported in the main text. The rest of the simulation parameters are described in table 1 of the main text.

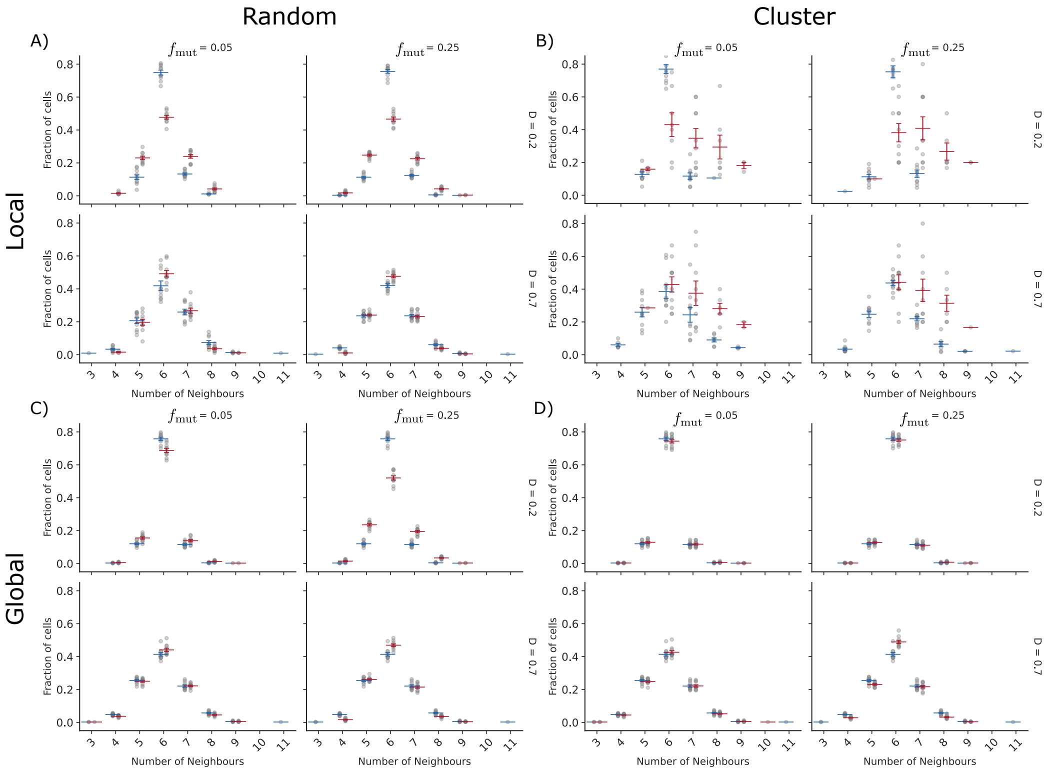


Figure S4: Distribution of polygon sides at $t = 0$ (blue markers) and after $T_2$-mediated mutant removal is complete, at $t = 15000$ (red markers), the same evaluation time as the heatmaps of the main text. Panels correspond to randomly distributed (A, C) and clustered (B, D) mutant configurations, shown for the local (A, B; mutant-neighbourhood wild-type cells, i.e. wild-type neighbours of mutant cells, updated while each mutant cell is present and fixed once it is cleared) and global (C, D; all cells) measurements. Within each panel, columns correspond to $f_{\mathrm{mut}} = 0.05$ and 0.25 and rows to $D = 0.2$ and $D = 0.7$, as indicated. Horizontal bars give the mean over $N_{\mathrm{sim}} = 10$ independent simulations and vertical bars the standard error of the mean; grey markers show the individual realisations. In the clustered configuration the local distribution shifts asymmetrically towards classes above six, with heptagons, octagons and nonagons all gaining at the expense of hexagons; this asymmetry is absent from both the random configurations and the corresponding global distributions. The rest of the simulation parameters are described in table 1 of the main text.
